# Temporal scaling in *C. elegans* larval development

**DOI:** 10.1101/2020.09.21.306423

**Authors:** Olga Filina, Rik Haagmans, Jeroen S. van Zon

## Abstract

It is essential that correct temporal order of cellular events is maintained during animal development. During post-embryonic development, the rate of development depends on external conditions, such as food availability, diet and temperature. How timing of cellular events is impacted when the rate of development is changed at the organism-level is not known. We used a novel time-lapse microscopy approach to simultaneously measure timing of oscillatory gene expression, hypodermal stem cell divisions and cuticle shedding in individual animals, during *C. elegans* larval development from hatching to adulthood. This revealed strong variability in timing between isogenic individuals under the same conditions. However, this variability obeyed ‘temporal scaling’, meaning that events occurred at the same time when measured relative to the duration of development in each individual. We also observed pervasive changes in population-averaged timing when temperature, diet or genotype were varied, but with larval development divided in ‘epochs’ that differed in how the timing of events was impacted. Yet, these variations in timing were still explained by temporal scaling when timing was rescaled by the duration of the respective epochs in each individual. Surprisingly, timing obeyed temporal scaling even in mutants lacking *lin-42/Period*, presumed a core regulator of timing of larval development, that exhibited strongly delayed, heterogeneous timing and growth arrest. Timing of larval development is likely controlled by timers based on protein degradation or protein oscillations, but such mechanisms do not inherently generate temporal scaling. Hence, our observations will put strong constraints on models to explain timing of larval development.

## Introduction

Numerous cellular events that occur during animal development, such as cell division, cell movement and gene expression, must be tightly coordinated in time to allow formation of a functional organism with a correctly established body plan. However, despite our increasing understanding of the regulation of developmental timing[1-3], how cells in developing organisms measure time and execute events in the correct temporal order remains poorly understood. Moreover, the rate of post-embryonic development in animals is significantly affected by external conditions, such as food availability, diet and temperature. For example, severe dietary restriction extends the duration of larval development in the nematode worm *C. elegans* as much as ten-fold, without resulting in apparent defects in development[4]. How the timing of individual developmental events is adjusted in response to such changes in the organism-level rate of development is not known.

This question about developmental timing has a parallel in the context of spatial patterning during development. It has been shown that spatial gene expression patterns often scale with organ or embryo size, i.e. with the spatial pattern adjusted in each individual organ or embryo so that the spatial features occurred at the same position relative to its overall size[5-8]. For example, in *Drosophila* embryos gap genes are expressed in bands along the anteroposterior body axis[9, 10]. These bands have highly stereotypical positions relative to the embryo’s size, even though this size shows significant variability between individuals[6]. Moreover, embryos of closely related species that vary greatly in size exhibit the same number of bands with similar position relative to the size of the embryo[6]. Here, we examine whether, analogous to scaling of spatial patterns in development, the timing of development exhibits temporal scaling, meaning that, when the organism-level rate of development is changed, the timing of individual events is adjusted so that they still occur at the same time when measured relative to the total duration of development. Such a mechanism would ensure the correct synchrony of developmental events even when organism-level timing is changed in an unpredictable manner by shifts in external conditions.

Due to its invariant cell lineage and highly stereotypical development. *C. elegans* is an ideal model organism to study developmental timing. Its postembryonic development consists of four larval stages (L1-L4) that are separated by a molting event, where a new cuticle is synthesized and the old cuticle shed[11]. After the final L4 molt, animals reach reproductive maturity, marking their transition into adulthood. There is a clear periodic aspect to *C. elegans* development, with molts occurring every 8-10 hours at 25°C. Moreover, larval stages are accompanied by genome-wide oscillatory expression of a multitude of genes, with peaks occurring once per larval stage[12-14].

Molecular mechanisms that have been proposed for regulation of developmental timing include oscillators, that encode time in periodic changes in protein level, and ‘hourglass’ mechanisms, that record time in the steady accumulation or degradation of proteins[15]. Developmental timing has been extensively studied in *C. elegans*, leading to the discovery of heterochronic genes[2, 3]. Heterochronic genes such as *lin-14* and *lin-28* show expression levels that decrease during larval development, suggestive of an hourglass mechanism[16, 17]. Indeed, mutations that perturb the *lin-14* and *lin-28* temporal expression patterns lead to timing defects, with events in one larval stage shifted to an earlier stage or repeated in subsequent stages[18]. At the same time, such mutations have only limited impact on developmental timing on the organism level. In contrast, the heterochronic gene *lin-42* is expressed in an oscillatory manner during development, peaking once every larval stage. In *lin-42* mutants, developmental timing is severely perturbed, with strong animal-to-animal variability in larval stage duration[19]. The body-wide, oscillatory expression dynamics of *lin-42,* together with its impact on larval stage duration, makes *lin-42* an interesting candidate for a global regulator of developmental timing. Intriguingly, *lin-42* is a homolog of Period, an important component of the circadian clock in *Drosophila* and higher organisms[20]. Hence, it has been speculated that *lin-42* forms part of an oscillator-based timer that allows cells and organs to read out developmental time[11].

How timing of individual events is impacted by changes in the organism-level rate of development is poorly characterized. Timing of *C. elegans* larval development is often measured at the population level, by examining the developmental stage of animals sampled from age-synchronized populations. This approach has limited time resolution and does not allow measuring timing of multiple events within the same individual. The latter is a particular problem for mutants such as *lin-42,* where developmental synchrony between individual animals is lost. However, the alternative approach of following individual animals was so far performed manually, limiting the number of animals that could be examined. We have recently developed a novel microscopy approach that allows automated imaging of individual *C. elegans* larvae during their entire development and at single-cell resolution[21], making it possible to measure timing of cellular events in many individual larvae. Here, we used this approach to simultaneously measure the timing of three recurring developmental events (oscillatory expression of a molting cycle gene, hypodermal stem cell divisions and cuticle shedding) in individual *C. elegans* larvae, both under changes in environmental conditions (temperature and diet) and in mutant animals, that increased the duration of larval development up to ~3-fold.

Our measurements uncovered extensive variability in event timing between individuals, even in isogenic animals under identical environmental conditions. Strikingly, this variability obeyed temporal scaling, meaning that events occurred at the same time when rescaled by the total duration of development in each individual. Moreover, changes in average timing between populations that differ in genotype or environmental conditions could not be explained by a simple change in the overall rate of development of the organism. Instead, we found larval development is divided into distinct epochs, which are sequences of events that, upon variation in conditions or genotype, exhibit changes in timing that are identical, and differ from changes in timing observed for events in other epochs. Yet, this variation in timing between populations is also obeys temporal scaling, provided that event times were rescaled by the duration of individual epochs, rather than total duration of development. Surprisingly, this was even the case for a *lin-42* deletion mutant that showed strongly perturbed and highly variable developmental timing, suggesting that while *lin-42* is crucial for setting the duration of larval stages, it is dispensable for controlling event timing relative to each larval stage.

Overall, our results show that the broad variation observed between individuals, environmental conditions and genotypes in the timing of cellular events during *C. elegans* post-embryonic development can be fully captured by the simple concept of temporal scaling, thereby revealing a precise adaptation of cell-level timing to changes in the organism-level rate of development. These observations raise the important question how temporal scaling is implemented by the molecular mechanisms that control timing of larval development.

## Results

To examine how developmental timing is coordinated at the organism-level in developing *C. elegans* larvae, we measured the timing of multiple developmental events that occurred frequently and throughout all of larval development, focusing on seam cell divisions, oscillatory gene expression and ecdysis. Seam cells are hypodermal stem cells that divide asymmetrically once every larval stage, yielding one hypodermal cell and one seam cell (Fig. 1A,D)[22]. In addition, the seam cells V1-V4 and V6 undergo an additional symmetric division in the L2 larval stage, doubling their number. Oscillatory gene expression is a pervasive phenomenon in *C. elegans,* with thousands of gene transcripts oscillating during development with a periodicity corresponding to that of the molting cycle[12-14]. Here, we focused on the oscillatory gene *wrt-2,* a hedgehog-like protein expressed in seam cells, that peaks in expression once every larval stage[12, 21] (Fig. 1B,D). Finally, ecdysis is the shedding of the old cuticle at the end of each larval stage (Fig. 1C,D). By focusing on these three events, we captured qualitatively different developmental processes, while their repetitive nature allowed us to capture many events in a single experiment.

**Figure 1.**
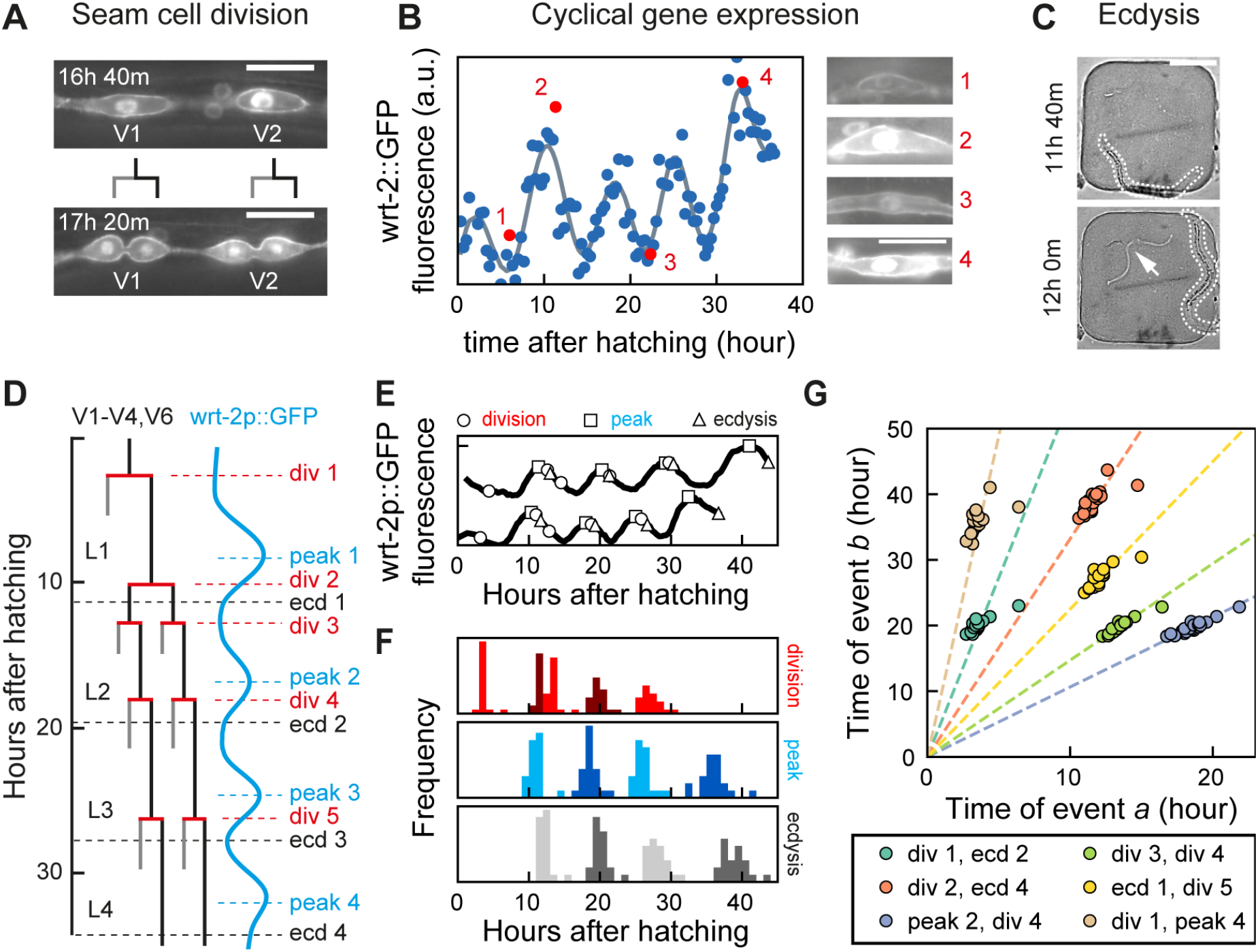
Scaling of developmental timing in individual animals. **(A)-(C)** Measuring timing of seam cell division, oscillatory gene expression and ecdysis. **(A)** Seam cells divide once or twice every larval stage giving rise either to hypodermal (grey line) or seam cell (black line) daughters. The timing of V1-V6 seam cell division was determined using the *wrt-2p::GFP::PH; wrt-2p::GFP::H2B* fluorescent reporter (*wrt-2p::GFP*). Scale bar: 15 μm. **(B)** Oscillatory *wrt-2* expression was visualized in seam cells of *wrt-2p::GFP* animals (panels 1-4, showing posterior V3 seam cell). We quantified *wrt-2p::GFP* fluorescence averaged over the V1-V6 seam cells (circles) and fitted the data to Eq. 1 (grey line) to determine the time of each *wrt-2* peak. Red circles are the time points in the side panels. Scale bar: 15 μm. **(C)** Time of ecdysis was determined by the appearance of a shed cuticle (arrow) away from the larva (outlined). Scale bar: 100 μm. **(D)** Schematic overview of seam cell division, *wrt-2* peak and ecdysis events during larval development. The V5 seam cell lineage differs by lacking the L2-specific symmetric division. **(E)** Example of variability in developmental timing between two wild-type animals. Markers indicate the timing of seam cell divisions (circles), wrt-2p::GFP fluorescence peaks (squares) and ecdyses (triangles). Tracks are shifted along the vertical axis for clarity. **(F)** Distribution of timing of seam cell divisions (red), wrt-2p::GFP fluorescence peaks (blue) and ecdyses (grey), measured for standard conditions, wild-type animals fed *E. coli* OP50 diet at 23°C (n=21). **(G)** Measured times of event pairs *a, b* for animals in (F). Each marker is an event pair measured in a single animal. Times of event pairs show temporal scaling, i.e. they lie clustered along lines of constant 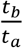 even as individual event times *t_a_* and *t_b_* show significant variability between individuals. Dashed lines are fits *t_b_* = *s_a,b_* · *t_a_* to the data for each event pair.

To accurately measure timing of individual events, we used a novel fluorescence time-lapse microscopy approach to follow the full ~40 h of post-embryonic development of individual *C. elegans* larvae with single-cell resolution[21]. Briefly, embryos were placed inside hydrogel chambers filled with *E. coli* as a food source. These chambers contained sufficient food to sustain development into adulthood, while constraining animals to the field of view of the microscope at each stage. By capturing fluorescence and transmitted light images with fast, 1-10 ms, exposure time, we could image developmental dynamics in individual cells inside moving larvae, without immobilizing animals.

To visualize seam cell divisions, we used the strain *heIs63[wrt-2p::H2B::GFP,wrt-2p::PH::GFP]*, where green fluorescent protein (GFP) was targeted specifically to the nucleus and membrane of seam cells[23]. Since divisions of individual seam cells occurred close together in time, we defined the time of each round of divisions as the average time at which V1-V6 cells have divided or started dividing, as determined by formation of the metaphase plate. Because the reporter used to detect seam cell divisions (*heIs63*) produces GFP under control of the *wrt-2* promoter, it enabled simultaneous measurement of oscillatory *wrt-2* expression. Fluorescent images were analyzed automatically using custom-written software to extract the mean fluorescent GFP intensity in seam cell nuclei at every time frame. Finally, the time of expression peaks was extracted from the resulting oscillatory *wrt-2* expression profiles, by fitting their dynamics with a combination of Gaussian functions and a linear offset (Fig. 1B, Eq. 1 in Methods).

Finally, the time of the ecdysis was defined as the time when the shed cuticle was first visible in the transmitted light image (Fig. 1C). For full details on data acquisition and analysis, see Methods.

### Scaling of developmental timing in individual animals

We first quantified timing of seam cell division, *wrt-2* expression peaks and ecdysis under standard conditions: wild-type animals fed *E. coli* OP50 at 23°C (Fig. 1E,F). Individual animals showed strong variability in the total duration of development (~40 h), defined as the time between hatching and L4 ecdysis, with a ~10 h difference observed between the first and last animal to enter adulthood. We observed similar variability in the timing of all measured developmental events (Fig. 1F). For some events, such as the second *wrt-2* peak and fourth seam cell division, the magnitude of the variability was larger than the average difference in timing, raising the question how the correct order of events was maintained. Interestingly, this variability in timing was strongly correlated: when we plotted event times *t_a_* and *t_b_* measured in the same animal against each other, for different pairs of events *a, b,* all data points clustered along a line (Fig. 1G). This strong correlation was even present for pairs of events that are widely separated in time, e.g. the L1 seam cell division and the L4 *wrt-2* peak.

A simple argument could explain this observation. As measure of developmental progression, we first defined the phase 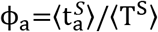, where 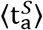 and 〈T^S^〉 are the population-averaged time of event *a* and duration of development under standard conditions (23°C), with *ϕ* = 0 and 1 corresponding to the start of larval development and adulthood. If we now assumed that for any event *a* the event time *t_a_*, measured in an individual animal, scales with the total duration of development *T* for that animal, *t_a_* = *ϕ_a_* · *T*, even as the duration *T* varies significantly between individuals, then the time at which two events *a* and *b* occur within same individual is related by 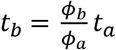, independent of total duration of development *T* in that individual. As a result, measurements for individual animals will be clustered along a line of constant 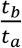, as observed experimentally. Strikingly, this observation holds for all pairs of events we examined (Supplementary Fig. 1), meaning that if an animal executed its first seam cell division earlier than the rest of the population, it was highly likely to be similarly early in executing all subsequent events for the rest of larval development, thereby ensuring the correct order of events despite the observed variability. We could reproduce these experimental results with a simple stochastic timing model (Supplementary Fig. 2, Eq. 2-3), if we assumed that the observed variability in timing was dominated by animal-to-animal variation in the organism-level rate of development, 1/*T*, that otherwise remained constant throughout development. Overall, our results show that the measured changes in timing between individuals can be fully explained by simple rescaling with the total duration of development in each individual.

### Scaling of developmental timing upon changes in temperature

Changes in environmental conditions typically increase or decrease the population-averaged duration of development, yet how timing of cell-level events is adjusted to such changes in organism-level timing is an open question. To address this, we first measured event timing in individual animals maintained at different temperatures, as the duration of larval development increases with decreasing temperatures[24]. As expected, we observed that as temperature was reduced from the standard temperature of 23°C to 19°C and 15°C, the duration of larval development increased from 39±2 to 57±1 and 105±2 hours, respectively (Supplementary Fig. 3). Likewise, we found that individual events were delayed more strongly with decreasing temperature (Fig. 2A).

**Figure 2.**
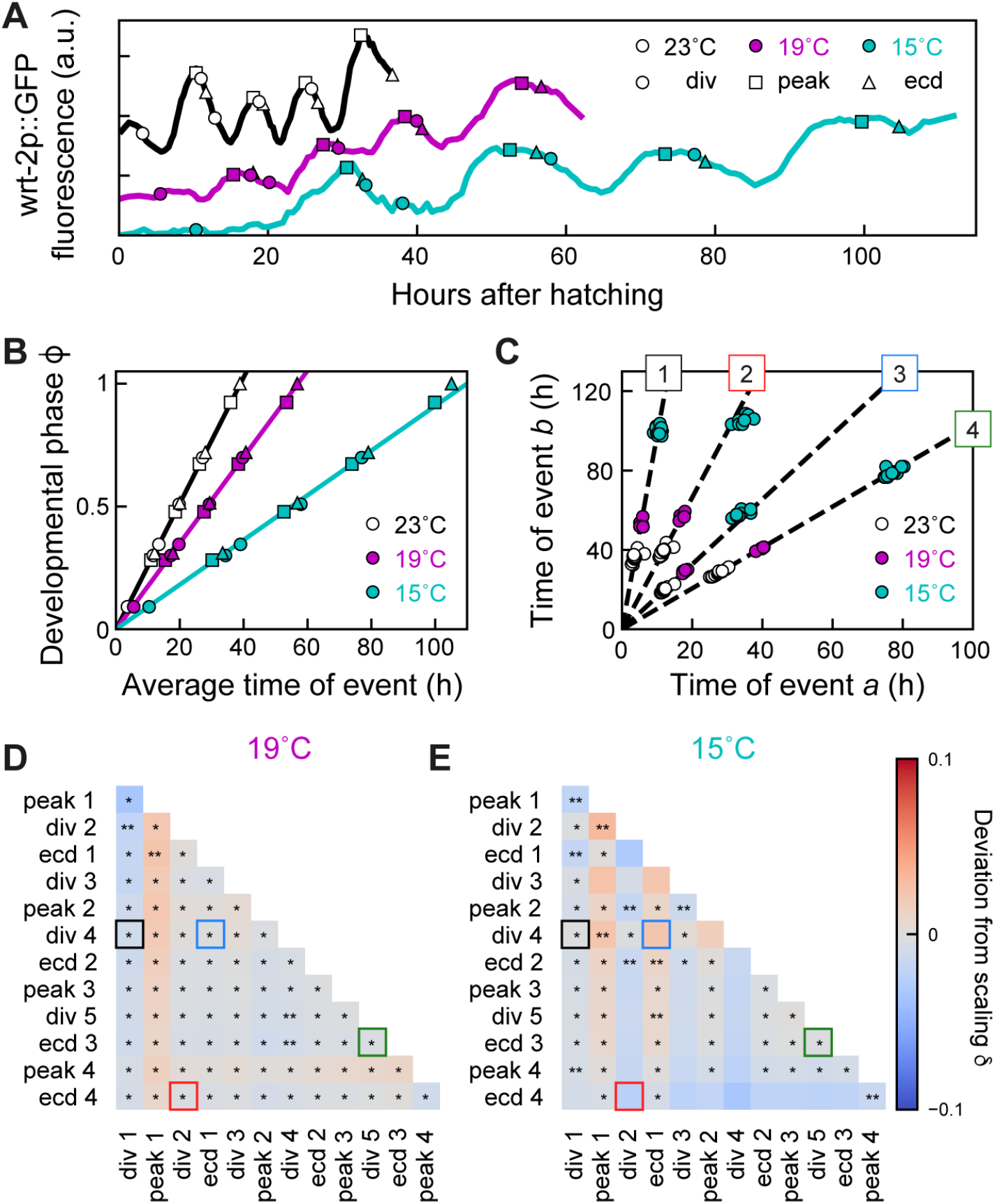
Temporal scaling of developmental timing at different temperatures. **(A)** Examples of developmental timing in individual animals at standard conditions (23°C, black), 19°C (magenta) and 15°C (cyan). Markers indicate seam cell divisions (circles), wrt-2p::GFP peaks (squares) and ecdyses (triangles). **(B)** Developmental phase as function of time for different temperatures. Developmental progression is modeled as the evolution of a developmental phase *ϕ* in time. Each developmental event *a* occurs at a specific phase 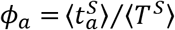, where 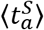 and 〈*T^S^*〉 are the population-averaged time of event *a* and total duration of development under standard conditions. With this definition, the phase for standard conditions (black line) increases linearly with rate 1/〈*T^S^*〉. Markers indicate the subsequent events as in (a). For 19°C or 15°C, the phase increases linearly with the average measured event times, albeit at lower rate compared to 23°C, with its evolution well fitted by the ‘Uniform’ model (magenta and cyan lines, defined in Fig. 3). **(C)** Measured times for different event pairs: (1) division 1 and 4, (2) division 2 and ecdysis 4, (3) ecdysis 1 and division 4, and (4) division 5 and ecdysis 3. Lines are a linear fit to data for standard conditions (23°C). For each event pair, data measured at different temperatures cluster along the same line. This means all events occurred at the same time relative to the total duration of development, despite strong variation in this duration both between individuals and conditions. **(D),(E)** Deviation from scaling *δ* for all event pairs for development at (D) 19°C and (E) 15°C. In addition, stars indicate the probability that data for standard conditions and 19°C or 15°C observe the same scaling relation, as assessed by a Kolmogorov-Smirnov (K-S) test, *:N.S., **:P<0.01, and P<0.001 otherwise. For animals at 19°C, temporal scaling was observed for all event pairs, while some significant deviations were seen at 15°C. Colored squares correspond to event pairs in (C).

To examine the impact of changing temperatures on average event timing, we examined the time evolution of the developmental phase *ϕ*. Our earlier definition ensured that under standard conditions (23°C) *ϕ* increases with constant rate 1/〈T^S^〉 (Fig. 2B). If the total duration of development increases or decreases, e.g. due to shifting environmental conditions, *ϕ*(*t*) will change, so that the same developmental phase *ϕ* is reached at a different time *t* compared to standard conditions. When we measured the average time of each seam cell division, wrt-2p::GFP peak and ecdysis for 19°C and 15°C, we found that, while the phase indeed increased at a lower rate compared to 23°C, its increase was still linear in time (Fig. 2B), meaning that all measured differences in average timing were explained by changes in the organism-level rate of development, 1/〈T〉.

When we examined events times in individuals, we found similar variability in timing for animals at 19°C and 15°C compared to 23°C. Moreover, times of event pairs still clustered along lines, meaning that temporal scaling with inter-individual variability in the duration of development occurred also for these conditions (Fig. 2C). Strikingly, event pairs clustered along the exact same line independent of temperature. This is a manifestation of population-level temporal scaling, i.e. for all three temperatures events occurred at the same relative time, when rescaled with the population-averaged duration of development observed for each temperature, even as development at 15°C was slowed ~3-fold compared to 23°C.

We could reproduce these observations with a simple, phenomenological model (‘Uniform’ model, Fig. 3). We assumed that population-averaged changes in timing resulted from a uniformly lowered rate of development, as observed experimentally (Fig. 2B, 3A), and included both an animal-to-animal variation in this rate, that remained constant throughout development, as well as variability in the phase of each individual event (Eq. 3). Stochastic simulations showed that when variability was dominated by the variation in developmental rate, the times of event pairs were positioned along a line, reflecting variability in timing between individuals (Fig. 3B-D). Significantly, this line appeared the same as for the corresponding events under standard conditions. To test this for both model and experiments, we defined the deviation of scaling *δ* as the signed angle between the lines that fit the data for standard and perturbed conditions (Fig. 3C), with *δ*>0 meaning that the line for perturbed conditions has a higher slope than for standard conditions. Here, we used the difference between angles rather than slopes, as the slope diverges when the first event *a* occurs close to hatching, *t_a_*≈ 0. Model calculations showed that for the ‘Uniform’ model the deviation *δ*=0 for all event pairs (Eqs. 6,9), meaning that they indeed lie along the line predicted for standard conditions.

**Figure 3.**
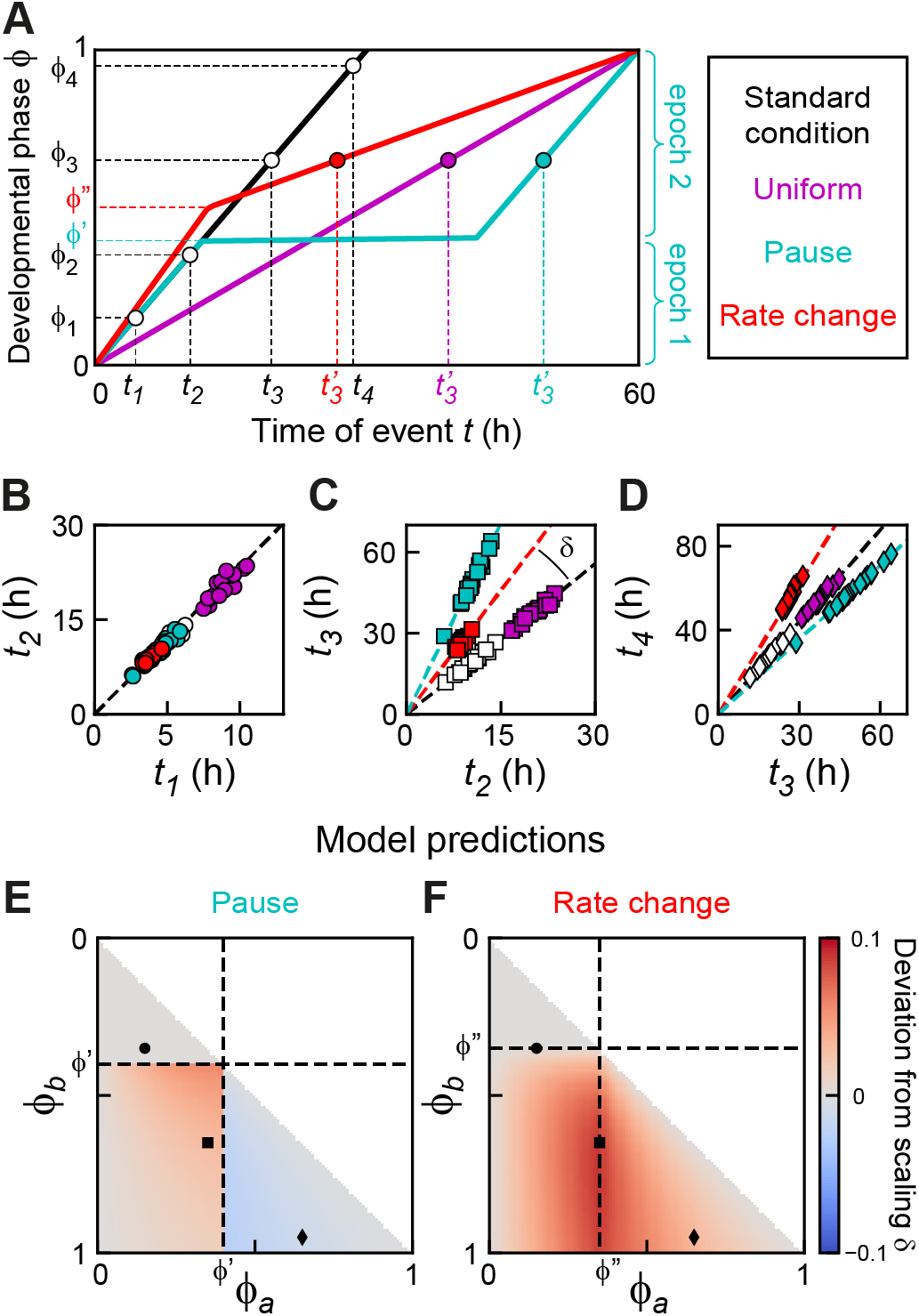
Timing models. **(A)** Developmental phase *ϕ* as function of time for three qualitatively different models that generate the same increase in duration of development, either by a uniform lower rate of development (magenta), a single, discrete pause at *ϕ*′ (cyan) or a single, discrete change in the rate of development at *ϕ*″ (red) (Eqs. 4-5 in Methods, parameters chosen for clarity). For the ‘Pause’ and ‘Rate change’ models, the discontinuity in developmental rate *dϕ/dt* separates development into two epochs of constant developmental rate, as illustrated for the ‘Pause’ model. **(B)-(D)** Simulated event times for the different event pairs indicated in (A) (Eq. 3). Marker color corresponds to the timing models in (A). For all models, times of event pairs are clustered along a line, i.e. occur at the same time when rescaled by each individual’s duration of development. For the ‘Uniform’ model, times of event pairs lie along the scaling line for standard conditions (dashed line), meaning that they occur at the same time when rescaled by the population-averaged duration of development. However, the other models often deviate from this scaling line. The deviation from scaling *δ* is defined as the signed angle between these two lines, as indicated in (C). **(E)-(F)** Predicted deviation from scaling *δ* for different events pairs *a* and *b* for the ‘Pause’ (E) and ‘Rate change’ (F) model (Eqs. 7-9), based on experimentally measured parameters (See Methods for parameters). Black markers correspond to the event pairs in (B)-(D). These results show that the timing models in (A) can be distinguished by measuring the deviation from population-level scaling and provide a quantitative prediction for the magnitude of the deviation that can be compared directly with experiments.

Next, we tested whether this held for our experimental observations. For each event pair *a* and *b* measured in the same individual, we calculated the angle 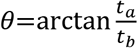 (See Methods for details) and the deviation as *δ* = 〈*θ^P^*〉-〈*θ^S^*〉, the difference between the average angle for standard (*S*, 23°C) and perturbed (*P*, 19°C or 15°C) conditions, with *δ*≈0 indicating that the data for standard and perturbed conditions clustered along the same line. In addition, we also used the two-sample Kolmogorov-Smirnov test to estimate the probability that the *θ* distributions measured for standard and perturbed conditions were draw from the same distribution. This analysis showed that most event pairs at 15°C and 19°C (Fig. 2D,E) lie along the same line as data for 23°C, i.e. changes in timing between temperatures are fully captured by temporal scaling with duration of development.

### Breakdown of population-level scaling upon changes in diet

To test whether temporal scaling is also observed under qualitatively different changes in environmental conditions, we studied the impact of changes in food uptake and diet on timing of individual events. Total duration of development can be changed by providing animals with other food than *E. coli* OP50[25-27]. Here, we used two different approaches. To mimic reduced food uptake, we fed the standard diet, E. coli OP50, to *eat-2(ad1113)* mutants that exhibit a 5-fold decrease in pharyngeal pumping and hence ingest bacteria at lower rate[28]. In addition, we grew wild-type animals on a diet of E. coli HB101, which was reported to have faster larval development[25].

The total duration of development was slightly different in *eat-2* mutants (40±2h) and wild-type animals on HB101 (38±1h), compared to standard conditions of wild-type animals on OP50 (39±2h, Fig. 4A, Supplementary Fig. 3). However, we observed more complex changes in timing when we examined the average timing of seam cell divisions, *wrt-2* peaks and ecdyses (Fig. 4B). For *eat-2* mutants, the developmental phase increased linearly with time, with a lower rate compared to standard conditions, consistent with the ‘Uniform’ model. However, for animals fed HB101 the increase of phase in time could not be fit by a single constant rate. Instead, we found that development separated into multiple epochs, sequences of events that different in how their timing was impacted by changing diet: events in the first epoch (hatching to third seam cell division) occurred with the same timing as under standard conditions, while timing in the second epoch (second *wrt-2* peak to L3 ecdysis) was best fit with the phase increasing at the same rate, but with a ~2h delay compared to standard conditions. We constructed a phenomenological model analogous to the ‘Uniform’ model, that incorporated the pause separating both epochs (‘Pause’ model, Fig. 3A). When we performed stochastic simulations of the ‘Pause’ model (Figs. 3B-D, Eqs. 3-4), we found that the times of simulated event pairs still clustered along a line, reflecting that animal-to-animal variability still obeyed temporal scaling, but one that differed from the equivalent line for standard conditions, meaning that differences in timing were not fully explained by rescaling with the population-averaged duration of development. Indeed, despite significant variability in timing between individuals, both for *eat-1* mutants and wild-type animals fed HB101, event pairs still clustered along a line (Fig. 4C), indicating that event times scaled with interindividual variability in developmental duration. Interestingly, while for *eat-1* mutants these data points were positioned along the same line as standard conditions, in animals fed HB101 we found that for some event pairs most points exhibited small, but systematic deviations from this line (e.g. event pairs 2 and 3 in Fig. 4C), as predicted by the ‘Pause’ model.

**Figure 4.**
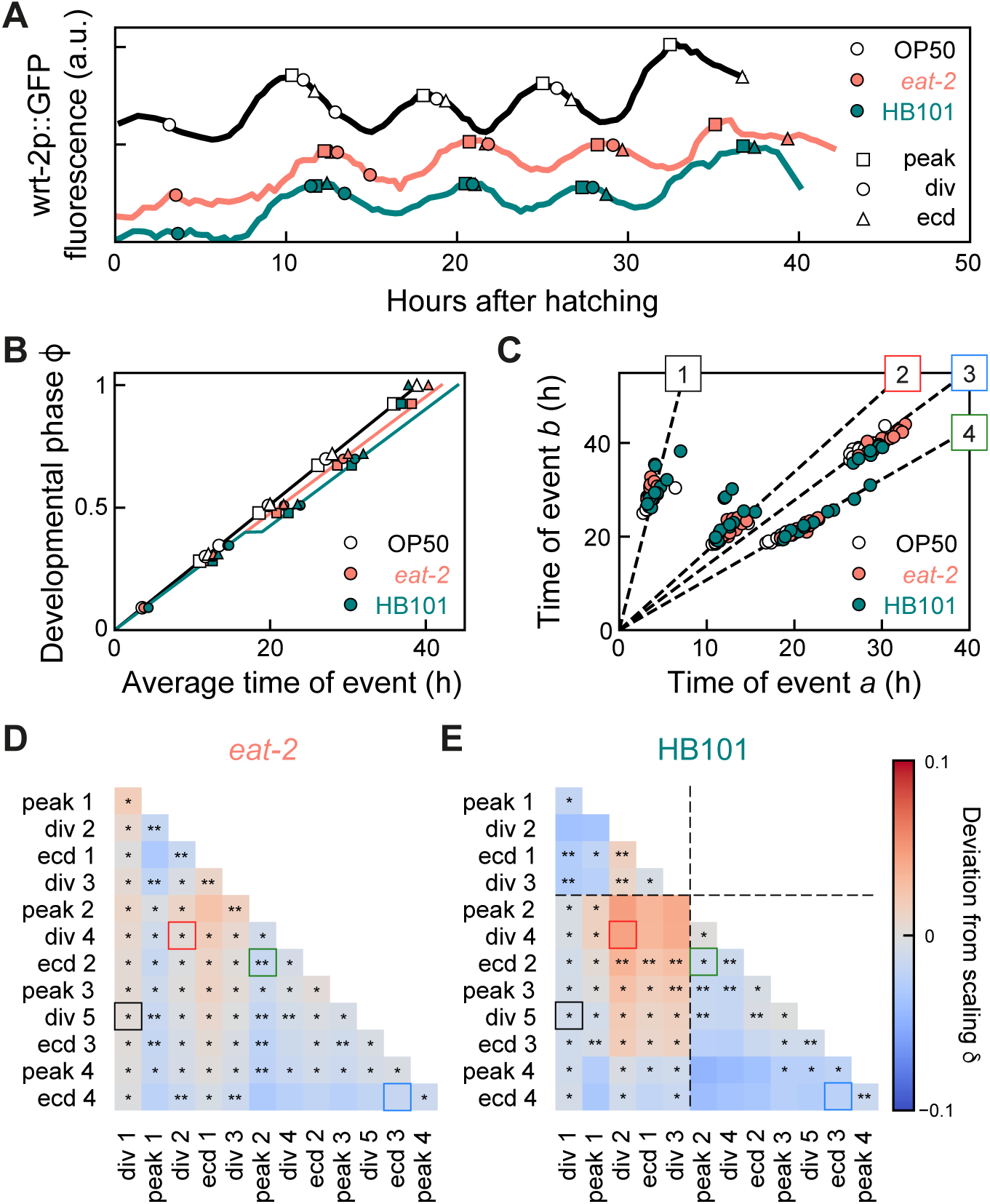
Breakdown of temporal scaling for change in diet. **(A)** Developmental timing in individual animals in standard conditions (black: wild-type, *E. coli* OP50 diet) and two conditions that exhibit slow development: wild-type on *E. coli* HB101 (green) and *eat-2* mutants, that have reduced food uptake, on *E. coli* OP50 (orange). Markers indicate seam cell divisions (circles), wrt-2p::GFP peaks (squares) and ecdyses (triangles). **(B)** Developmental phase as function of time. Markers indicate events as in (A). For *eat-2* mutants, the phase evolution is well fitted by the ‘Uniform’ model (orange line), and for wild-type on HB101 by the ‘Pause’ model (green line) with a ~2 h pause between division 3 and peak 2. **(C)** Measured times for event pairs: (1) division 1 and 5, (2) division 2 and 4, (3) peak 2 and ecdysis 2, and (4) ecdysis 3 and 4. Lines are a fit to data for standard conditions. For (2) and (3), animals on HB101 show systematic deviations from scaling. **(D),(E)** Deviation from scaling for (D) *eat-2* mutants and (E) animals fed HB101. Stars indicate the probability that the data observes the same scaling relation as on standard conditions: *:N.S., **:P<0.01, and P<0.001 otherwise (K-S test). Dashed lines in (E) indicate the time of the pause. While scaling is observed for almost all event pairs in *eat-2* mutants, animals on HB101 exhibit systematic deviations from scaling that match those predicted by the ‘Pause’ model both in sign and magnitude.

To directly compare these deviations between model and experiments, we calculated the deviation from scaling *δ* for the ‘Pause’ model, with the three parameters (the rate of development, and phase and duration of the pause) constrained by our experimental measurements (Fig. 4B, see Method for parameter values). Interestingly, the model predicted that these deviations occurred in a specific pattern (Fig. 3E): while timing of event pairs that both occurred before the developmental pause matched the line for standard conditions, deviations occurred when at least on event occurred after the pause. Event times clustered along a line with higher slope (*δ* > 0) when one event occurred before and the other after the pause, with stronger deviations if both events were close to the pause. In contrast, when both events occurred after the pause, event times clustered along lines with lower slope (*δ* < 0), with deviations stronger when events occurred farther apart in development. We then tested these predictions from the ‘Pause’ model by calculating *δ* for all measured event pairs. As predicted, we found a clear difference between *eat-2* mutants on OP50 and wild-type animals on HB101 (Fig. 4D,E). Whereas data for *eat-2* mutants largely clustered along the lines predicted by data for standard conditions, the data for animals fed HB101 showed significant deviations. Interestingly, these deviations strongly resembled, both in magnitude and sign, those predicted by the ‘Pause’ model, with a positive deviation when one event occurs before and the other after the pause, and a negative deviation when both occur after. This quantitative agreement implied that all deviations from scaling were due to the discontinuity in developmental rate that separated the two epochs, and that events occurred at the same time within each epoch when rescaled by the duration of that epoch in each individual.

Overall, these results show that changes in diet (HB101 instead of OP50) impact developmental timing in a manner that is more complex than rescaling with the population-averaged duration of development. However, the model suggests a simple picture that explains the experimental data: the deviations from population-level temporal scaling resulted from the pause at the mid-L2 larval stage found for animals fed HB101, with scaling observed within the two epochs separated by this pause. Moreover, each animal executes this temporal evolution of the developmental phase at an intrinsic speed that varies between individuals, giving rise to inter-individual temporal scaling for all event pairs.

### Perturbed developmental timing and growth in *lin-42* mutants

For wild-type animals on HB101 we observed deviations from temporal scaling even as diet had only minor impact on total duration of development. To seek stronger perturbations of developmental timing and, potentially, temporal scaling, we measured event timing in mutants of the heterochronic gene *lin-42*, for the following reasons. First, *lin-42* plays an important role in molting, with mutants showing longer larval stages, strongly reduced synchrony between individuals in progression through larval stages and frequent developmental arrest, with all these phenotypes increasing in severity as development proceeds[19, 29, 30]. Second, *lin-42* mutants exhibit heterochronic phenotypes in multiple organs[19, 20, 31, 32], indicating a body-wide role for *lin-42.* In addition, *lin-42* is expressed in many cells throughout the body and in a striking oscillatory manner, with LIN-42 protein levels peaking once every larval stage[20]. This, together with the homology of *lin-42* to the circadian clock gene period, has led to the speculation that *lin-42* acts as a global developmental timer[11, 19]. Finally, *lin-42* regulates the expression of many miRNAs, including those involved in timing through the heterochronic pathway and binds to the promoter of many genes[33-35]. Because of this wide-ranging impact on developmental timing and gene expression, *lin-42* appeared a prime candidate also for a core component of a potential scaling mechanism. Hence, we wanted to test whether *lin-42(0)* animals displayed stronger deviations from temporal scaling than observed under changes in diet.

We used the *lin-42(ox461)* allele that deletes the entire *lin-42* locus and shows the strongest perturbation of molting cycle progression[29]. Indeed, we found that postembryonic development at 23°C was significantly slower in *lin-42(0)* animals, with much stronger variability both in the total duration of development (57±7h for *lin-42(0),* compared to 39±2h for wild-type animals) and the duration of individual larval stages (Supplementary Fig. 3). In addition, *lin-42(0)* animals showed reduced growth, as measured by the increase of body length over time in individual animals (Fig. 5A,B). In particular, we observed a fraction of animals that stopped growing completely between the L2 and L4 larval stage, with some reaching body lengths of only 0.3mm, compared to 0.9mm for wild-type animals. Surprisingly, all animals that arrested growth appeared to otherwise continue development: they underwent multiple rounds of ecdysis, seam cell divisions and *wrt-2* expression peaks (Fig. 5C). Animals frequently skipped the final seam cell division and ecdysis, a heterochronic phenotype observed before[19], but did so independently of the growth arrest. After molting, *lin-42(0)* animals often remain stuck in their old cuticle, and it was suggested that this interferes with the ability to feed[29]. However, we observed growth arrest also in animals that appear to shed their cuticle normally. Moreover, growth-arrested animals also displayed pharyngeal pumping, suggesting that growth arrest was not simply caused by inability to take up food.

**Figure 5.**
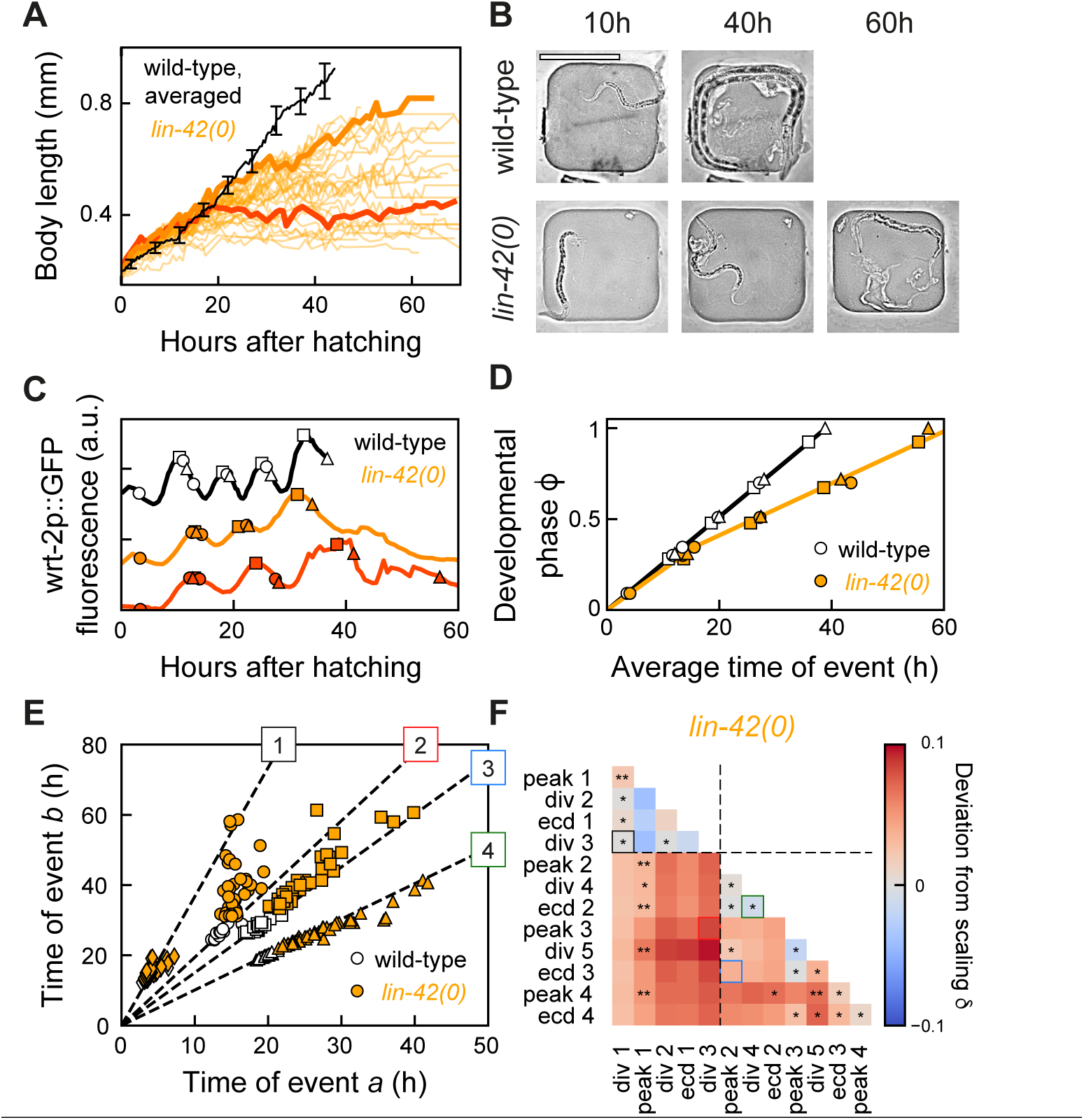
Growth arrest and temporal scaling in *lin-42(0)* animals. **(A)** Body length as function of time for standard conditions (black, average for wild-type animals at 23°C) and *lin-42(0)* animals (orange lines, individual animals). While some *lin-42(0)* animals only exhibited reduced growth, other animals showed complete growth arrest from the mid-L2 stage, ~20 h after hatching, capping body length to <0.5mm. Thick lines highlight one individual with reduced growth (light orange) and one arresting growth (dark orange). Error bars indicate S.E.M. **(B)** Microscopy images of a wild-type (upper panels) and growth-arrested *lin-42(0)* larva (lower panels) at different times after hatching. Scale bar: 200 μm. **(C)** Developmental timing in a wild-type larva (black), and in a slow-growing (light orange) and growth-arrested (dark orange) *lin-42(0)* animal that correspond to the individuals highlighted in (A). Markers indicate seam cell divisions (circles), wrt-2p::GFP peaks (squares) and ecdyses (triangles). The growth-arrested animal (dark orange) executed most events that occur after mid-L2, indicating the absence of a developmental arrest. **(D)** Developmental phase as function of time in wild-type (black) and *lin-42(0)* animals (orange). The phase evolution in *lin-42(0)* animals was well fitted with the ‘Rate change’ model (orange line), occurring at wild-type rate until the mid-L2 stage, and ~2-fold decreased rate for events occurring later. **(E)** Measured times for event pairs: (1) division 1 and 3, (2) division 3 and peak 3, (3) peak 2 and ecdysis 3, and (4) division 4 and ecdysis 2. Lines are a fit to data for standard conditions. Overall, *lin-42(0)* animals show strong variability in timing between individuals, and for event pairs (2) and (3) exhibit clear deviations from scaling. **(F)** Deviation from scaling for development in *lin-42(0)* mutants. Stars indicate probability that *lin-42(0)* data observes the same scaling relation as wild-type: *:N.S., **:P<0.01, and P<0.001 otherwise (K-S test). Deviations from scaling resemble those predicted by the ‘Rate change’ model, with dashed lines indicating the measured time of the rate change.

### Breakdown of inter-individual and population-level temporal scaling in *lin-42* mutants

When we compared the average timing of each seam cell division, *wrt-2* peak and ecdysis between wild-type and *lin-42(0)* animals at 23°C (Fig. 5D), we found that again development separated into multiple epochs that differed in impact on timing: in the first epoch (hatching to third seam cell division) developmental phase in increased at the same rate in *lin-42(0)* and wild-type animals, while in the second epoch (from the second *wrt-2* peak onwards) the phase still increased linearly in *lin-42(0)* mutants but with a strongly decreased rate compared to wild-type animals. When we examined timing in individual animals, we found that variability in timing was dramatically increased in *lin-42(0)* mutants. Yet, despite this strong variability data points for the majority event pairs still clustered along a line (Fig. 5E, Supplementary Fig. 4). Strikingly, we found that this was not the case for all event pairs: for instance, some animals that were among the first to execute the third seam cell division exhibited an exceptionally late third *wrt-2* peak, resulting in many points away from the line (Fig. 5E, event pair 2), indicating a breakdown of inter-individual temporal scaling. The key assumption underlying inter-individual temporal scaling is that the variation in the developmental rate between individuals remains constant throughout larval development. However, in individual *lin-42(0)* animals, the duration of early-larval development was often poorly correlated with the duration of late-larval development, compared to the other genotypes and conditions we examined (Supplementary Fig. 5A-C). Indeed, model simulations showed that this observation was sufficient to explain these deviations from inter-individual scaling (Supplementary Fig. 5D-G).

For event pairs that did cluster along a line, data points often showed systematic deviations from the line predicted by data for standard conditions (wild-type animals at 23°C) corresponding to a lack of population-level temporal scaling. We used the measured evolution of developmental phase to construct a phenomenological model of timing in *lin-42(0)* mutants (‘Rate change’ model, Fig. 3A, 5D). This model reproduced the experimental observation that data for most event pairs clustered along lines, but with systematic deviations from the lines predicted by data for standard conditions (Fig. 3B-D). In particular, for the ‘Rate change’ model these lines always had a larger slope than for standard conditions, i.e. *δ* ≥ 0 (Fig. 3F). Indeed, when we calculated *δ* for all experimentally measured event pairs (Fig. 5F) the resulting dependence of *δ* on the developmental phase strongly resembled the prediction from the ‘Rate change’ model (Fig. 3F): first, the deviation was largest for event pairs with one event occurring prior to the mid-L2 larval stage, when we observed the change in developmental rate, and the other occurring afterwards. Second, in contrast to data for wild-type animals fed HB101, the deviation in *lin-42(0)* mutants was always positive (*δ* > 0). These results suggested that the breakdown of population-level temporal scaling was largely accounted for by the change in developmental rate at the mid-L2 larval stage that separated the two observed epochs. Finally, compared to the rest of the population, growth-arrested animals did not develop more slowly, and their developmental timing did not exhibit stronger deviations from temporal scaling (Supplementary Fig. 4), suggesting this was largely independent of physical growth.

Overall, *lin-42(0)* mutants exhibited strongly perturbed and variable timing, with larval development separated into two epochs that differed in developmental rate. Surprisingly, given the putative role of *lin-42* as a key regulator of developmental timing, we found that both individual variability and population-level changes in timing were explained by temporal scaling, but only when considering events within the same epoch, i.e. events occurred at the same time when measured relative to the duration of epochs in each individual.

## Discussion

Here, we showed that changes in timing of individual developmental events, under a broad array of conditions that change the total duration of *C. elegans* larval development, are largely explained by temporal scaling, both for variability in timing between individuals and for population-level changes in timing when environmental conditions or genotype are varied.

We found that many of our experimental observations could be reproduced by simple, phenomenological timing models (Fig. 3). In these models, the complexity of developmental progression is reduced to the evolution of a developmental phase in time, similar to the use of phase in the analysis of nonlinear oscillators. Animal-to-animal variability arises because each animal proceeds through its phase evolution at an intrinsically different rate, giving rise to the strongly correlated variability we measured for timing of event pairs. When we compared event times between animals of different genotype or raised under different conditions, the resulting changes in timing were not simply explained by the population-level change in total duration of larval development. We found that these deviations from scaling were explained by the subdivision of larval development in distinct epochs, that differ in how the evolution of phase is impacted upon changes in genotype or environmental condition. This result implies that event timing did exhibit temporal scaling, but only when taking into account these epochs: events occur at the same time, independent of genotypes or conditions, when measured relative to the duration of each epoch rather than the total duration of development. While these phenomenological models do not provide a molecular mechanism for temporal scaling, they reveal a remarkably simple organization that unifies the broad variations in timing seen in our experiments.

It is striking that isogenic animals under identical environmental conditions showed strong variability in developmental timing. This variability in timing was strongly correlated between event pairs, for almost all event pairs, conditions and genotypes examined (Fig. 1G, 2C, 4C, 5E). This implies that each individual progresses through development at a constant rate, which however varies from animal to animal, with this variability in overall developmental rate explaining the observed variability in timing. What controls this variation in developmental rate? A recent study found that animal-to-animal variability in timing of the L4 ecdysis, i.e. the transition into adulthood, correlated with age of the mother at fertilization[36], with embryos generated by older mothers developing more rapidly. This was linked to the amount of yolk proteins loaded in each embryo, which increased with the mother’s age, suggesting that the rate of larval development is determined by the status of nutrient stores in the embryo. It is surprising, however, that this variation in rate persists throughout development, particularly as larvae depend for growth on feeding rather than internal stores. The only case we observed where the rate of early-larval development was not predictive of the late-larval rate in *lin-42(0)* mutants (Supplementary Fig. 5). This might reflect a role of *lin-42* in maintaining the developmental rate at the embryo-set level, but could also reflect that the severity of the *lin-42(0)* phenotype, as it emerges during larval development, is independent of the early-larval developmental rate.

For some changes in genotype or conditions (changes in temperature, *eat-2* feeding mutants), the rate of development was reduced by a constant factor for all larval development (Figs. 2B, 4B). In contrast, both for wild-type animals fed HB101 and *lin-42(0)* mutants, we observed a discontinuity in the rate of development that separated larval development into two epochs (Figs. 4B, 5D). Interestingly, even though the nature of the discontinuity differed, with a pause for animals fed HB101 and a change in rate for *lin-42(0)* mutants, both occurred at the same development stage: the mid-L2 larval stage, between third seam cell division and second *wrt-2* peak. This stage is notable, as it forms an important transition both in terms of development and metabolism. First, under conditions of crowding or food deprivation larvae enter dauer, an alternative developmental state that is highly stress-resistant, and commit to this fate before the mid-L2 larval stage[37, 38]. Interestingly, *lin-42* is involved in dauer commitment, with *lin-42* mutants showing increased dauer formation at 27°C[39]. Hence, the change in developmental rate at this stage might reflect involvement of the dauer decision-making machinery. However, we prefer another explanation: *C. elegans* larvae also shift their metabolism between the L1 and L2 larval stage, from the glyoxylate cycle to the TCA cycle[40]. The glyoxylate cycle likely allows L1 larvae to use stored lipids as an energy source, potentially rendering their development less dependent on ingestion of food. As a consequence, shifts in diet (from *E. coli* OP50 to HB1010) or inability to ingest or metabolize food in *lin-42(0)* mutants might only impact developmental timing substantially after shifting to the TCA cycle upon entering the L2 larval stage.

Our observations raise the question how temporal scaling is regulated. One attractive mechanism to regulate timing in a manner that is synchronized throughout the body and adapts to changes in rate of development under different conditions, is to couple developmental timing to physical growth. If timing was regulated in such a way that each event occurred at a specific body size, it would explain temporal scaling, as changes in conditions that impact the body’s growth rate would naturally lead to concomitant changes in developmental timing. Analogously, cell cycle timing in bacteria and yeast strongly depends on cell size and growth[41-44]. Indeed, progression of *C. elegans* larval development is tightly linked to body size[24], and under dietary conditions that caused slow growth, larval stages lengthened so that molts occurred at their stereotypical body size[4]. Moreover, conditions that do not allow growth, such as starvation, typically lead to developmental arrest at the start of each larval stage[45-47]. It is therefore an important observation that *lin-42(0)* mutants continued development without physical growth (Fig. 4), and with timing of development obeying temporal scaling within each epoch (Fig. 5). Continued development after growth arrest in *lin-42(0)* mutants was shown recently for motor neuron, gonad and vulva development[29], but timing was not examined. Our work now extends these observations, indicating that continued development without growth is likely a *lin-42(0)* phenotype with body-wide impact, and specifically invalidates a mechanism where temporal scaling results from coupling developmental timing to physical growth.

Two distinct classes of molecular mechanisms to explain temporal scaling could be envisioned. In the first class, feedback mechanisms or checkpoints actively adjust timing. An example of this class are the size-based checkpoints proposed for single-celled organisms, but invalidated by us as a mechanism for *C. elegans*. A more general example are the feedback mechanisms that actively shape morphogen gradients underpin that underlie spatial scaling in developmental pattern formation[8]. The second class of mechanisms emerged from recent studies examining the dependence of developmental timing in *C. elegans*, fly and frog embryos on temperature[48-51]. They explain temporal scaling as emerging from careful tuning of the molecular processes in the cell. Specifically, Kurtz and Eisen found that timing of fruit fly development scaled uniformly with temperature[50], similar to our observations for *C. elegans* larval development (Fig. 2). Intriguingly, in these studies the measured dependence of timing on temperature followed an Arrhenius equation[48-50], which is historically used to describe the temperature dependence of chemical reactions. Temporal scaling arises only if timing of all events follows the same Arrhenius equation, i.e. timing of all events varies by the same factor for a given change in temperature. Because the rates of molecular processes are likely not only controlled by temperature but also by metabolite levels, such thermodynamic mechanisms might also generate temporal scaling upon changes in diet or food uptake, as we observed (Fig. 4,5). However, more recent measurements disputed the key assumption that all processes follow the same Arrhenius equation[48, 49], raising the question whether thermodynamic mechanisms by themselves are sufficient to explain temporal scaling, or if additional active feedback or checkpoint mechanisms are still essential.

One of the most enduring mysteries of development is how its timing is regulated. Whereas we have a deep understanding of how spatial patterns arise during development[8], our understanding of how events like cell division, cell movement or gene expression are controlled in time is still very limited. Post-embryonic development poses a particular challenge, as its rate of progression depends strongly on environmental conditions such as food availability. How timing is adapted on the cellular level in response to such organism-level changes is an open question. On the molecular level, developmental timing is thought to be controlled either by oscillators or ‘hourglass’ mechanisms. In *C. elegans,* even though there is now evidence both for hourglass mechanisms[16, 17] and oscillators[11, 14] in regulating timing of larval development, we still do not understand how the exact time at which cell-level events are initiated is determined on the molecular level. Both hourglass and oscillator mechanisms do not exhibit temporal scaling of their decay time or period, respectively, without alterations to their basic mechanism. Hence, our observation of temporal scaling will put strong constraints on possible models that explain developmental timing of *C. elegans* larval development.

## Materials and methods

### Strains and experimental conditions

The following strains were used: *heIs63[Pwrt-2::GFP::PH;Pwrt-2::GFP::H2B;Plin-48::mCherry]*[23], *eat-2(ad1113)*[28], *lin-42(ox461)*[29]. To monitor *wrt-2::GFP* expression in *eat2* and *lin-42* mutants, these mutations were crossed into *heIs63*. The *eat-2* genotype was confirmed by measuring the rate of pharyngeal pumping, which is decreased 5-fold compared to wild-type animals[28]. As *lin-42* constitutes a complex genetic locus encoding multiple isoforms, we chose to use the *lin-42(ox461)* allele that deletes the entire locus of *lin-42[29].* In addition, the *lin-42(ok2385)* allele that deletes the main isoform *lin-42a* and the overlapping exons of *lin-42b,* was analyzed and showed similar phenotypes as *lin-42(ox461)* (data not shown). For maintenance, all strains were cultured at 20°C on NGM (Nematode growth medium) agar plates seeded with OP50 strain of *E. coli* bacteria, using standard *C. elegans* techniques. For time-lapse experiments, we refer to standard conditions as *wrt2p::GFP* animals fed *E. coli* OP50 at 23°C. For experiments in perturbed conditions we varied one experimental parameter (*C. elegans* strain, temperature or diet) while keeping the others unchanged. For the diet experiment in Fig. 4, *E. coli* HB101 was used instead. In that case, animals were maintained on HB101 for 5-7 generations prior to the experiment.

### Time-lapse microscopy

Custom time-lapse microscopy setup was used to monitor the entire development of individual *C. elegans.* Late-stage embryos were placed inside the 250×250×20 μm polyacrylamide microchambers (one embryo per chamber) filled with *E. coli* bacteria as food source. Nikon Ti-E inverted microscope with a large chip camera (Hamamatsu sCMOS Orca v2) and a 40x magnification objective (Nikon CFI Plan Fluor 40x, NA=1.3, oil immersion) were used for imaging. Transmission and fluorescence images were acquired with an LED light source (CoolLED pE-100 615nm) and a 488 nm laser (Coherent OBIS LS 488-100), respectively. Each chamber containing a single animal was imaged every 20-40 minutes during the entire larval development (40-100 hours depending on the genotype and temperature). A stack of 20-30 images in the Z-direction was acquired using short exposure times (1-10 ms), such that the motion of the animal was insignificant. We have previously demonstrated that this technique does not hinder larval growth and development[21].

### Temperature control

All experiments were performed in a temperature-controlled room with a constant temperature inside the sample of 23°C. To perform an experiment at different temperature, an additional temperature control system was used. A thermoelectric chiller (Thermotek T257P) was used to cool the custom made objective jacket by circulating an antifreeze fluid (a mixture of water and glycerin) between the chiller and the objective jacket. In order to calibrate the system, a thermocouple temperature sensor measuring 0.025 mm in diameter (RS Pro) was placed inside the sample in contact with the polyacrylamide hydrogel and connected to a digital thermometer (RS Pro). The temperature was then varied on the thermoelectric chiller while the resulting temperature inside the sample was being monitored. In this work, experiments were performed at 23, 19 or 15 °C.

### Image analysis

Time-lapse image stacks were processed with custom Python software in order to obtain the precise timing of ecdysis events, seam cell divisions, and peaks in oscillatory *wrt-2* expression. For every animal, the times of hatching and ecdysis events were annotated based on visual inspection of transmitted light images. Hatching was defined as the time larvae first appears out of the egg shell, while ecdysis events were defined as the first appearance of the shed cuticle in the chamber (Fig. 1C). Times of seam cell divisions were annotated based on visual inspection of the *wrt-2p::GFP* fluorescence signal in the nucleus and the membrane of seam cells. Divisions of V1-V6 seam cells occur close together in time. We therefore defined the time of each round of divisions as the average time V1-V6 cells had divided or had started dividing, as determined by the formation of the metaphase plate. We only analyzed seam cells located on the side of the body closest to the objective.

To determine the time of peaks in oscillatory *wrt-2* expression, we first obtained *wrt-2* expression profiles as a function of time for individual animals. For this, in every time frame we automatically segmented the region encompassing seam cells using a Watershed algorithm and calculated the average fluorescence intensity inside this region. Finally, to find the time of each peak (*μ_i_*), we fitted the obtained oscillatory profiles with a combination of Gaussian functions and a linear offset using non-linear least-squares minimization (Fig. 1B):

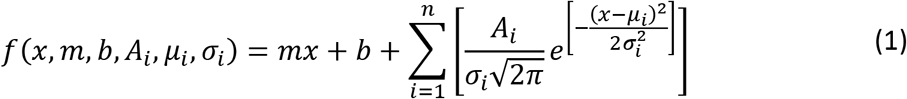

where *n* is the number of peaks, *A_i_, μ_i_, ρ_i_* are the amplitude, center and width of peak. Finally, we fitted the experimentally measured times for pairs of events *a* and *b* to a line function of the form *t_b_ = s_a,b_·t_a_* using non-linear least-squares minimization (Fig. 1G), using the Linear Model from the *lmfit* package in Python. To measure the animal’s body length as a function of time (Fig. 5A), we manually annotated ~10 points along the anterior-posterior body axis and performed spline interpolation. Body length was defined as the length of the resulting spline curve.

### Timing models and simulations

We model the progression of development as the evolution of a developmental phase *ϕ*, that increases from *ϕ*=0 (start of larval development at hatching) to *ϕ*=1 (entry into adulthood at the L4 ecdysis). The exact assignment of a phase to a particular developmental event is arbitrary. Here, we define the phase so that for standard conditions (wild-type animals at 23 °C) the phase increases linearly with time, 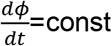, and *ϕ*=1 at time *t=T*, where *T* is the total duration of development at standard conditions. As a result, for standard conditions we use the following definition for the developmental phase of event *a*:

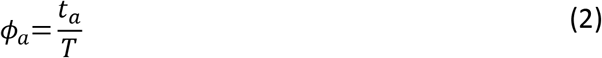

For other conditions or genotypes, the time evolution *ϕ*(*t*) of the developmental phase has a different form. As a result, the time of event *a*, occurring at a developmental phase *ϕ_a_* is given by *t_a_ = f*(*ϕ_a_*), where *f* is a monotonically increasing function that is specific for each condition or genotype. Expressions for *f*(*ϕ*) are discussed further below.

To incorporate animal-to-animal variability, we assumed two different sources of variability. First, there is an intrinsic variability in the stage *ϕ_a_* at which each event *a* occurs, that is uncorrelated between different events occurring within the same animal. Second, we assumed variability in the total duration of development, *T*. This corresponds to an animal-to-animal variability in the global rate of development that impacts each event occurring within the same animal equally. Then, the event time *t_a,i_* for event *a* in animal *i* is given by:

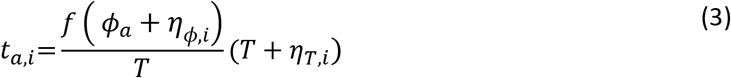

where *T* and *ϕ_a_* correspond to the population average values, while *η_T_* and *η_ϕ_* are Gaussian noise sources with standard deviation *σ_T_* and *σ_ϕ_*, respectively.

The function *f*(*ϕ*) changes for differing environmental conditions or mutants that perturb the duration of development. In particular, we considered three different models, the ‘Uniform’, ‘Pause’, and ‘Rate change’ models (Fig. 3A). For the ‘Uniform’ model, event times are given by Eq. 2, but now with an increased duration of development *T*’. For the ‘Pause’ model, developmental occurs at the same rate as for standard conditions, but with a pause at developmental phase *ϕ*′ that results in a total duration of development *T*′=(1 + *κ*)*T*, resulting in:

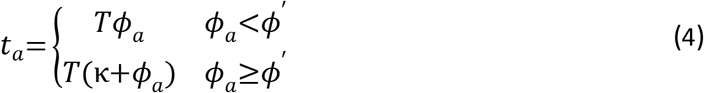

Finally, for the ‘Rate change’ the developmental rate differs between events occurring prior to a developmental phase *ϕ*” and events occurring afterwards, resulting in:

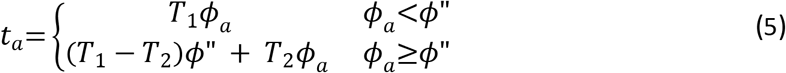

where 1/*T*_1_ and 1/*T*_2_ correspond to the two developmental rates and the total duration of development is given by *T*′=*T*_1_*ϕ*”+*T*_2_(1 − *ϕ*”).

### Calculation of deviation from scaling for timing models

For the ‘Uniform’, ‘Pause’ and ‘Rate change’ model, we calculate the deviation from scaling as follows. First, we use that fact that for events *a* and *b* that occur in the same animal, the total duration of development, *T’*, has the same value, to express *t_b_* as function of *t_a_*. For the ‘Uniform’ model, this yields:

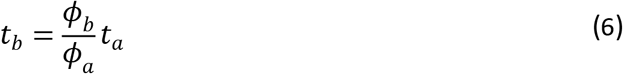

meaning that that event pairs lie along the same line as measured for individuals under standard conditions, and that the changes in timing can be fully captured by a simple rescaling of event times with the duration of development *T′* under non-standard conditions. In contrast, for the ‘Pause’ model, this yields:

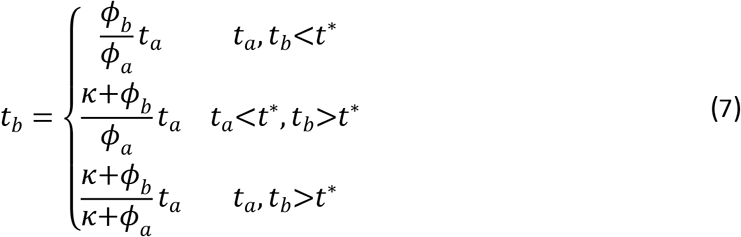

Here, pairs of time points for events *a* and *b* only lie along the same line as those for standard conditions when both events occur before the time of the delay, *t*\*=*Tϕ*′. Otherwise, the slope of the line is different from wild-type conditions and depends explicitly on the delay parameter *κ*. Finally, for ‘Rate change’ model, corresponding to the *lin-42(ox461)* mutant, we have:

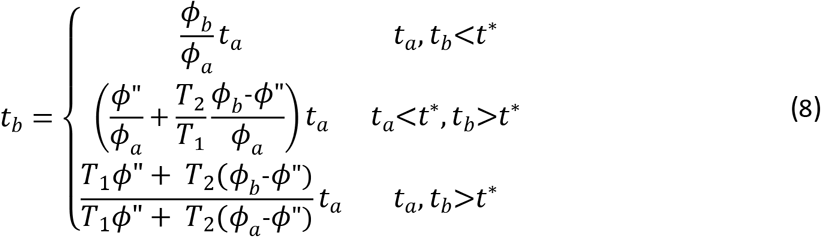

where *t*\* = *T*_1_*ϕ*”. Apart from the case when *t_a_, t_b_<t*\*, this expression depends explicitly on the parameters *ϕ*” and *T*_2_ and does not lie along the same line event pairs for standard conditions. Finally, we calculate the deviation from scaling as:

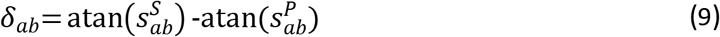

where *t_b_ = s_ab_ · t_a_*, with the slope *s_ab_* given by Eqs. 7-9, and *S* and *P* denote standard and perturbed conditions, respectively.

### Model parameters

For the model results in Fig. 3A-D, we used the following parameters, chosen to emphasize and clarify the differences between models. For the ‘Standard condition’ model: *T*=30 h. For the ‘Uniform’ model: *T*’=60 h. For the ‘Pause’ model: *κ*=1 and *ϕ*’=0.4, resulting in a total duration of development of *T*’=60 h. For the ‘Rate Change’ model: *T*’=25 h, *T*_2_=95 h and *ϕ*”=0.5, also resulting in *T*’=60 h. For the stochastic simulations in Fig. 3C-D: *σ_T_*=5 h and *σ_ϕ_*=7·10^−3^. For Figs. 2, 3E,F, 4 and 5, we used model parameters that were fitted to the experimental data. In Fig. 2, we used the following fitted parameter values: *T*^23°C^=39 h, *T*^19°C^=57 h, and *T*^15°C^=110 h. In Fig. 4 and Fig. 3E, we used the following fitted parameter values: *T*^*eat*−1^=42 h, *T*^HB101^=42 h, *κ*_HB101_=5·10^−2^ and 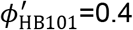. In Fig. 5 and Fig. 3F, we used the following fitted parameters: 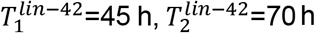 and 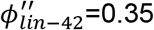. The parameters for HB101 and *lin-42* were used to make the quantitative predictions in Fig. 3E,F using Eqs. 7,8.

### Calculating the deviation from scaling for experimental data

We used the following procedure to assess whether the timing of event pairs *a* and *b* under perturbed conditions *P* deviate from the scaling relationship found for their timing under standard conditions *S*. For each animal *i* with measured event timing *t_a,i_* and *t_b,i_*, we calculate 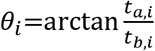. First, we calculate the average deviation *δ* from scaling as 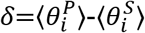. However, in particular for event pairs with small average event times 〈*t_a,i_*〉 and 〈*t_b,i_*〉, values of *θ_i_* can vary substantially, leading to non-zero deviation *δ* for the typical number of animals analyzed for these experiments. Hence, we also estimate the probability that the two series 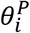 and 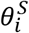 are sampled from the same distribution, using the two-sample Kolmogorov-Smirnov test (*ks_2samp* from the *scipy.stats* package in Python). We reported the P value, with high P meaning that the distributions of the two samples are likely the same, and, hence, obey the same temporal scaling relationship.

## Acknowledgements

Some strains were provided by the CGC, which is funded by NIH Office of Research Infrastructure Programs (P40 OD010440). We thank Sander van den Heuvel for providing the wrt-2p::GFP strain and Ann Rougvie for providing the *lin-42* alleles. We thank Pieter Rein ten Wolde and Tom Shimizu for feedback on the manuscript. This work is part of the research programme VIDI with project number 680-47-529, which is (partly) financed by the Dutch Research Council (NWO)

## Supplementary figures

**Supplementary Fig. 1.**
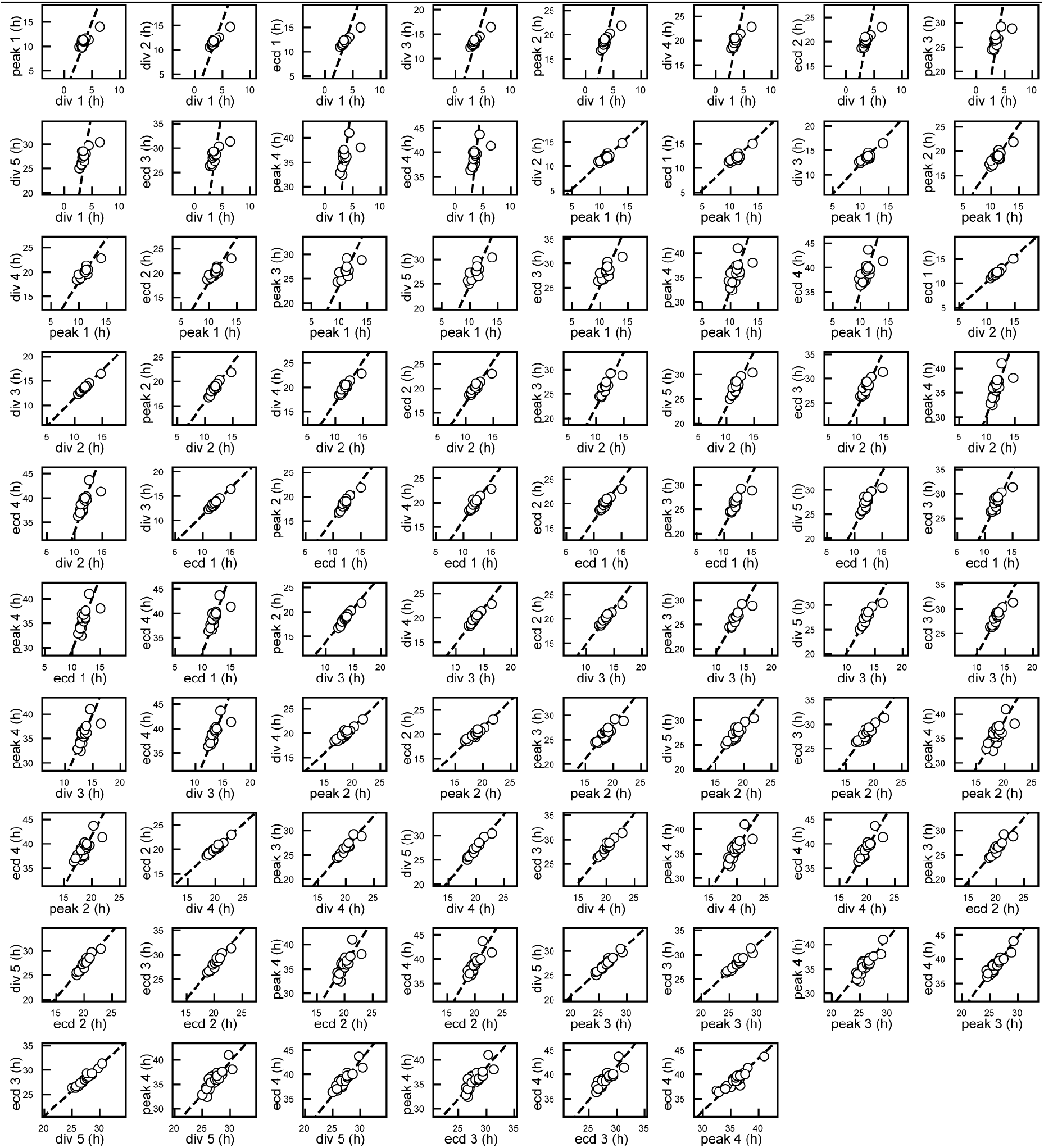
Temporal scaling in animal-to-animal variability of timing. Measured times of for all measured event pairs *a, b* in wild-type animals, on an *E. coli* OP50 diet at 23°C. Markers correspond to times measured in a single animal. Dashed lines are fits of the form *t_b_ = s_a,b_ · t_a_*.

**Supplementary Fig. 2.**
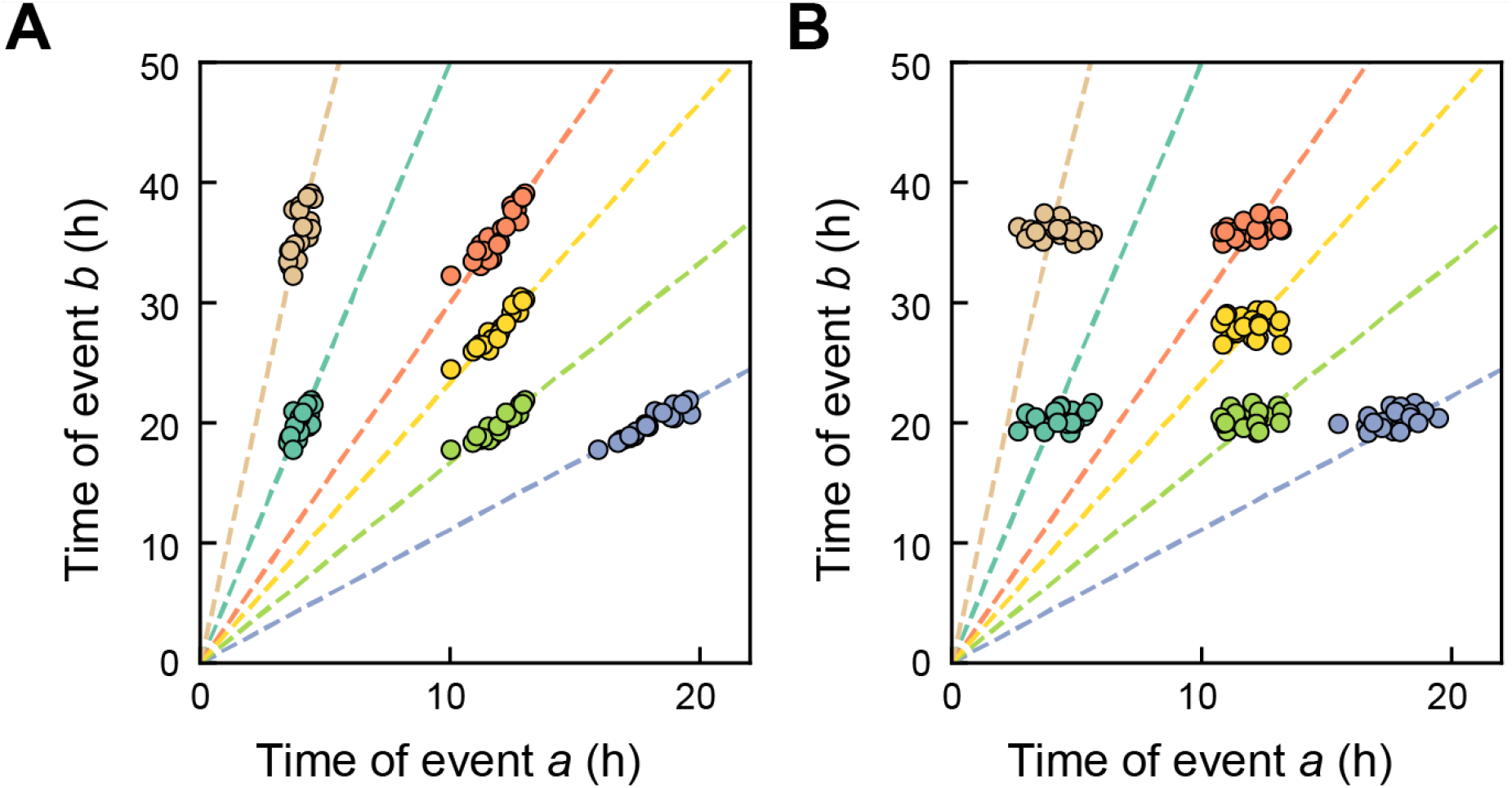
Stochastic timing model. **(A)** Simulated times of event pairs *a, b* in the stochastic timing model (Eqs. 2 and 3 in Methods). Simulation parameters were chosen to resemble the experimental data in Fig. 1e. Times were generated for six event that occur at developmental phases *ϕ*=0.1, 0.3, 0.45, 0.5, 0.7 and 0.9, that are chosen so that the plotted event pairs occur at similar times as those in Fig. 1G. The average duration of development was *T*=40h. For the noise sources, we used standard deviations *σ_T_*=3*h* and *σ_ϕ_*=7·10^−3^, meaning that common variability in the rate of development, 1/*T*, is stronger than variability in timing of each individual event. The simulated data closely resembles the experimental data in Fig. 1G, with times for event pairs *a, b* scattered along lines of 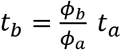 (dashed lines). **(B)** Same simulation as in (A), but now with standard deviations *σ_T_*=0.1*h* and *σ_ϕ_*=2·10^−2^, meaning that variability in the timing of individual events is stronger than that in the global rate of development. As a consequence, times for event pairs no longer cluster along lines of 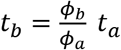 (dashed lines).

**Supplementary Fig. 3.**
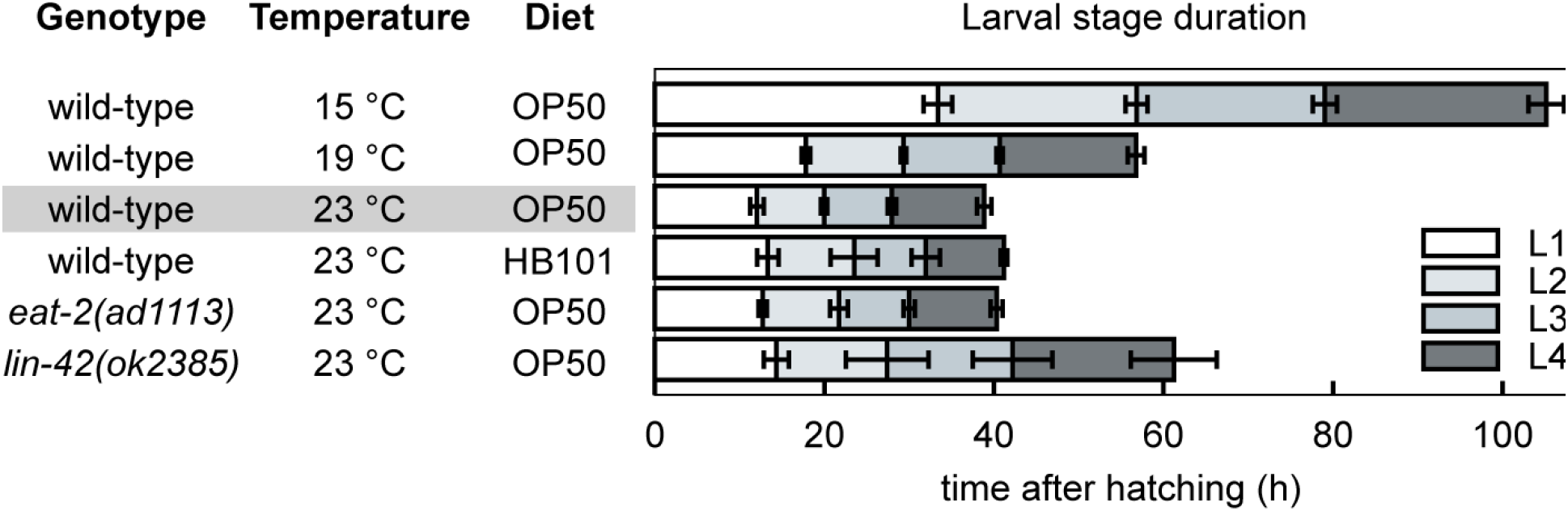
Overview of genotypes, conditions and duration of larval stages. Average duration of larval stages L1-L4 for the different genotypes and environmental conditions used. Wild-type refers to animals carrying only the *wrt-2p::GFP* reporter, which exhibit timing of larval stages similar to animals without this reporter. Strains with genotype different than wild-type also carry the *wrt-2p::GFP* reporter. OP50 and HB101 refer to different *E. coli* bacterial strains used a food source. The standard condition used as benchmark to compare event timing against is outlined in grey. For larval stage duration, error bars indicate S.D. and for each condition n>7, except for *lin-42(0)* mutants, where only a few animals complete the L4 stage (n=4).

**Supplementary Fig. 4.**
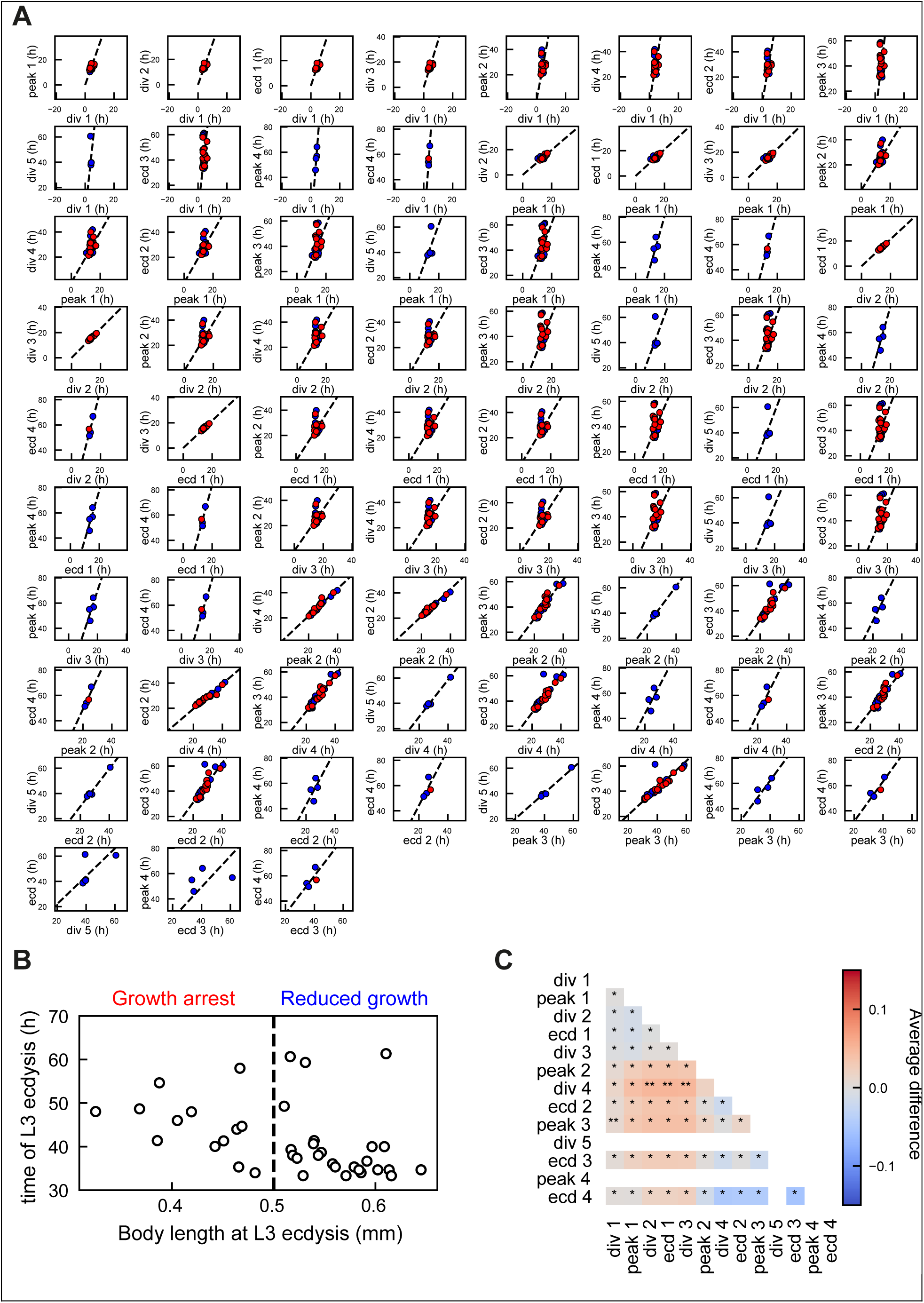
Dependence of deviation from temporal scaling on *lin-42(0)* growth arrest phenotype. **(A)** Measured times for all event pairs measured in at least *n>4* animals. Some event pairs found during wild-type L4 larval stage development do not appear as they are executed only rarely in *lin-42(0)* animals. Lines are a linear fit to the data points. Markers are colored according to growth phenotype, as defined in panel (B). The deviation from scaling does not differ strongly between growth-arrested animals (red) and animals with reduced growth (blue). **(B)** Length at L3 ecdysis compared to time of L3 ecdysis in *lin-42(0)* animals. Based on this, we separated the population in growth-arrested animals (length < 0.5mm at L3 ecdysis) and animals with reduced growth (as compared to wild-type animals). Growth-arrested animals developed more slowly than animals with reduced growth, but a small number of animals with reduced growth also displayed very slow development (L3 ecdysis later than 50 h after hatching). We scored the growth phenotype based on L3 characteristics, because most animals skip the L4 larval stage. **(C)** Difference in scaling between growth-arrested (GA) and reduce-growth (RG) animals. Color indicates the difference 〈*θ^GA^*〉 − 〈*θ^RG^*〉 between the two populations, where, for each event pair *a* and *b* measured in an individual animal, the angle 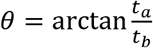. Stars indicate the probability that the distribution of *θ* is the same for growth-arrested and reduced-growth animals: *:N.S., **:P<0.01, and P<0.001 otherwise (K-S test). Overall, no significant differences in scaling were observed between growth-arrested and reduced-growth animals, indicating that growth-arrested animals do not display stronger breakdown of scaling. Empty squares reflect event pairs for which at least one of the two events did not occur in either growth-arrested or reduced-growth animals.

**Supplementary Fig. 5.**
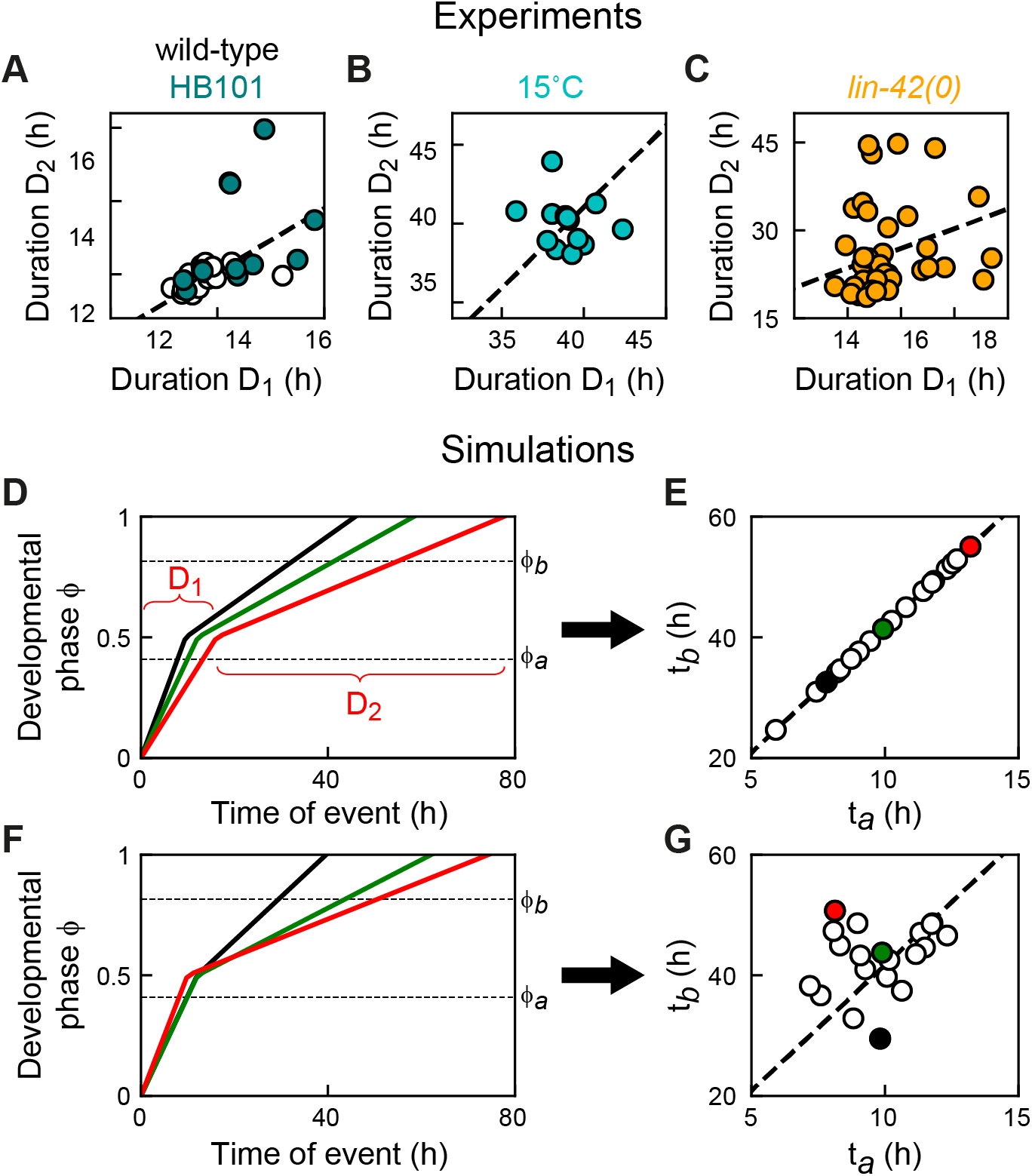
Breakdown of inter-individual scaling. **(A),(B),(C)** Experimentally measured correlation between the developmental duration *D*_1_ and *D*_2_ of development before and after the change in developmental rate, for development under standard conditions (A, wild-type at 23°C), animals fed HB101 (A), at 15°C (B) and in *lin-42(0)* mutants (C). The duration *D*_1_ is defined at the time from hatching to the third seam cell division, while *D*_2_ is given by the time from the third seam cell division to the third ecdysis. For *D*_2_, we do not include the time from the third to fourth ecdysis, as this last part of larval development is often skipped in *lin-42* mutants. The correlation between *D*_1_ and *D*_2_ is much weaker in *lin-42* mutants. **(D),(E)** Inter-individual variability in the ‘Rate change’ model. In this model, all individuals go through the same evolution of developmental phase *ϕ* but at a rate that differs between individuals (D). Hence, variability in the duration *D*_1_ and *D*_2_ of development before and after the change in developmental rate is strongly correlated within the same individual. As a consequence, times of events *t_a_* and *t_b_*, occurring at phase *ϕ_a_* and *ϕ_b_*, are clustered tightly along a line, indicating temporal scaling (E). The colored markers in (E) correspond to the individuals whose phase evolution is shown in (D). Model is given by Eqs. 3 and 5 in Methods, with *T*_1_=25 h, *T*_2_=95 h, *ϕ*”=0.5, *σ_ϕ_*=7·10^−3^ and *σ*_*T*_2__=15 h. Dashed line is Eq. 8. **(F),(G)** Breakdown of inter-individual scaling, when variability in durations *D*_1_ and *D*_2_ is not correlated within the same individual. As a result, individual animals show strong deviations from temporal scaling. Model is given by Eqs. 3 and 5, but *T*_1_ and *T*_2_ vary independently within the same individual, with *σ*_*T*_1__=3 h and *σ*_*T*_2__=15 h. This model result indicates that the lack of correlation between durations *D*_1_ and *D*_2_ observed in lin-42(0) mutants can explain the lack of inter-individual scaling seen in *lin-42(0)* mutants when comparing timing of events occurring before and after the change in developmental rate in Fig. 5E.

## Notes

### Competing Interest Statement

The authors have declared no competing interest.

### Summary of Updates

All figures and text have been revised for clarity

## References

1. Ebisuya, M. and J. Briscoe, What does time mean in development? 2018. 145(12).

2. Moss, E.G., Heterochronic genes and the nature of developmental time. Curr Biol, 2007. 17(11): p. R425–34.

3. Rougvie, A.E., Control of developmental timing in animals. Development, 2001. 2(9): p. 690–701.

4. Uppaluri, S. and C.P. Brangwynne, A size threshold governs Caenorhabditis elegans developmental progression. Proceedings of the Royal Society B: Biological Sciences, 2015. 282(1813): p. 20151283.

5. Ben-Zvi, D., et al., Scaling of the BMP activation gradient in Xenopus embryos. Nature, 2008. 453(7199): p. 1205–11.

6. Gregor, T., et al., Diffusion and scaling during early embryonic pattern formation. Proc Natl Acad Sci U S A, 2005. 102(51): p. 18403–7.

7. Houchmandzadeh, B., E. Wieschaus, and S. Leibler, Establishment of developmental precision and proportions in the early Drosophila embryo. Nature, 2002. 415(6873): p. 798–802.

8. Umulis, D.M. and H.G. Othmer, Mechanisms of scaling in pattern formation. Development, 2013. 140(24): p. 4830–43.

9. Driever, W. and C. Nüsslein-Volhard, A gradient of bicoid protein in Drosophila embryos. Cell, 1988. 54(1): p. 83–93.

10. Driever, W. and C. Nüsslein-Volhard, The bicoid protein determines position in the Drosophila embryo in a concentration-dependent manner. Cell, 1988. 54(1): p. 95–104.

11. Monsalve, G.C. and A.R. Frand, Toward a unified model of developmental timing: A “molting” approach. Worm, 2012. 1(4): p. 221–30.

12. Hendriks, G.J., et al., Extensive oscillatory gene expression during C. elegans larval development. Mol Syst Biol, 2014. 53(3): p. 380–92.

13. Kim, D., D. Grün, and A. van Oudenaarden, Dampening of expression oscillations by synchronous regulation of a microRNA and its target. Mol Cell, 2013. 45(11): p. 1337–44.

14. Meeuse, M.W. and Y.P. Hauser, Developmental function and state transitions of a gene expression oscillator in Caenorhabditis elegans. 2020. 16(7): p. e9498.

15. Rensing, L., U. Meyer-Grahle, and P. Ruoff, Biological timing and the clock metaphor: oscillatory and hourglass mechanisms. Chronobiol Int, 2001. 18(3): p. 329–69.

16. Moss, E.G., R.C. Lee, and V. Ambros, The cold shock domain protein LIN-28 controls developmental timing in C. elegans and is regulated by the lin-4 RNA. Cell, 1997. 88(5): p. 637–46.

17. Ruvkun, G. and J. Giusto, The Caenorhabditis elegans heterochronic gene lin-14 encodes a nuclear protein that forms a temporal developmental switch. Nature, 1989. 338(6213): p. 313–9.

18. Ambros, V. and H.R. Horvitz, Heterochronic mutants of the nematode Caenorhabditis elegans. Science, 1984. 226(4673): p. 409–16.

19. Monsalve, G.C., C. Van Buskirk, and A.R. Frand, LIN-42/PERIOD controls cyclical and developmental progression of C. elegans molts. Curr Biol, 2011. 21(24): p. 2033–45.

20. Jeon, M., et al., Similarity of the C. elegans developmental timing protein LIN-42 to circadian rhythm proteins. Science, 1999. 286(5442): p. 1141–6.

21. Gritti, N., et al., Long-term time-lapse microscopy of C. elegans post-embryonic development. Nat Commun, 2016. 7: p. 12500.

22. Sulston, J.E. and H.R. Horvitz, Post-embryonic cell lineages of the nematode, Caenorhabditis elegans. Dev Biol, 1977. 56(1): p. 110–56.

23. Wildwater, M., et al., Cell shape and Wnt signaling redundantly control the division axis of C. elegans epithelial stem cells. Development, 2011. 138(20): p. 4375–85.

24. Byerly, L., R.C. Cassada, and R.L. Russell, The life cycle of the nematode Caenorhabditis elegans. I. Wild-type growth and reproduction. Dev Biol, 1976. 51(1): p. 23–33.

25. Avery, L. and B.B. Shtonda, Food transport in the C. elegans pharynx. J Exp Biol, 2003. 206(Pt 14): p. 2441–57.

26. MacNeil, L.T., et al., Diet-induced developmental acceleration independent of TOR and insulin in C. elegans. Cell, 2013. 153(1): p. 240–52.

27. Soukas, A.A., et al., Rictor/TORC2 regulates fat metabolism, feeding, growth, and life span in Caenorhabditis elegans. Genes Dev, 2009. 23(4): p. 496–511.

28. Raizen, D.M., R.Y. Lee, and L. Avery, Interacting genes required for pharyngeal excitation by motor neuron MC in Caenorhabditis elegans. Genetics, 1995. 141(4): p. 1365–82.

29. Edelman, T.L., et al., Analysis of a lin-42/period Null Allele Implicates All Three Isoforms in Regulation of Caenorhabditis elegans Molting and Developmental Timing. G3 (Bethesda), 2016. 6(12): p. 4077–4086.

30. Olmedo, M., et al., A high-throughput method for the analysis of larval developmental phenotypes in Caenorhabditis elegans. Genetics, 2015. 201(2): p. 443–448.

31. Abrahante, J.E., E.A. Miller, and A.E. Rougvie, Identification of heterochronic mutants in Caenorhabditis elegans. Temporal misexpression of a collagen::green fluorescent protein fusion gene. Genetics, 1998. 149(3): p. 1335–51.

32. Tennessen, J.M., et al., Novel heterochronic functions of the Caenorhabditis elegans period-related protein LIN-42. Dev Biol, 2006. 289(1): p. 30–43.

33. McCulloch, K.A. and A.E. Rougvie, Caenorhabditis elegans period homolog lin-42 regulates the timing of heterochronic miRNA expression. Proc Natl Acad Sci U S A, 2014. 111(43): p. 15450–5.

34. Perales, R., et al., LIN-42, the Caenorhabditis elegans PERIOD homolog, negatively regulates microRNA transcription. PLoS Genet, 2014. 10(7): p. e1004486.

35. Van Wynsberghe, P.M. and A.E. Pasquinelli, Period homolog LIN-42 regulates miRNA transcription to impact developmental timing. Worm, 2014. 3(4): p. e974453.

36. Perez, M.F., et al., Maternal age generates phenotypic variation in Caenorhabditis elegans. Nature, 2017. 552(7683): p. 106–109.

37. Golden, J.W. and D.L. Riddle, The Caenorhabditis elegans dauer larva: developmental effects of pheromone, food, and temperature. Dev Biol, 1984. 102(2): p. 368–78.

38. Schaedel, O.N., et al., Hormonal signal amplification mediates environmental conditions during development and controls an irreversible commitment to adulthood. PLoS Biol, 2012. 10(4): p. e1001306.

39. Tennessen, J.M., K.J. Opperman, and A.E. Rougvie, The C. elegans developmental timing protein LIN-42 regulates diapause in response to environmental cues. Development, 2010. 137(20): p. 3501–11.

40. Wadsworth, W.G. and D.L. Riddle, Developmental regulation of energy metabolism in Caenorhabditis elegans. Dev Biol, 1989. 132(1): p. 167–73.

41. Cooper, S. and C.E. Helmstetter, Chromosome replication and the division cycle of Escherichia coli B/r. J Mol Biol, 1968. 31(3): p. 519–40.

42. Johnston, G.C., J.R. Pringle, and L.H. Hartwell, Coordination of growth with cell division in the yeast Saccharomyces cerevisiae. Exp Cell Res, 1977. 105(1): p. 79–98.

43. Mitchison, J.M. and J. Creanor, Further measurements of DNA synthesis and enzyme potential during cell cycle of fission yeast Schizosaccharomyces pombe. Exp Cell Res, 1971. 69(1): p. 244–7.

44. Robert, L., Size sensors in bacteria, cell cycle control, and size control. Front Microbiol, 2015. 6: p. 515.

45. Baugh, L.R., To grow or not to grow: nutritional control of development during Caenorhabditis elegans L1 arrest. Genetics, 2013. 194(3): p. 539–55.

46. Hu, P.J., Dauer. WormBook, 2007: p. 1–19.

47. Schindler, A.J., L.R. Baugh, and D.R. Sherwood, Identification of late larval stage developmental checkpoints in Caenorhabditis elegans regulated by insulin/IGF and steroid hormone signaling pathways. PLoS Genet, 2014. 10(6): p. e1004426.

48. Begasse, M.L., et al., Temperature Dependence of Cell Division Timing Accounts for a Shift in the Thermal Limits of C. elegans and C. briggsae. Cell Rep, 2015. 10(5): p. 647–653.

49. Crapse, J., et al., Evaluating the simple Arrhenius equation for the temperature dependence of complex developmental processes. 2020.

50. Kuntz, S.G. and M.B. Eisen, Drosophila embryogenesis scales uniformly across temperature in developmentally diverse species. PLoS Genet, 2014. 10(4): p. e1004293.

51. Chong, J., C. Amourda, and T.E. Saunders, Temporal development of Drosophila embryos is highly robust across a wide temperature range. J R Soc Interface, 2018. 15(144).

